# A Reference Tissue Implementation of Simultaneous Multifactor Bayesian Analysis (SiMBA) of PET Time Activity Curve Data

**DOI:** 10.1101/2024.12.04.626559

**Authors:** Granville J. Matheson, Johan Lundberg, Martin Gärde, Emma R. Veldman, Amane Tateno, Yoshiro Okubo, Mikael Tiger, R. Todd Ogden

**Author notes:** Shared last author (Mikael Tiger, R. Todd Ogden). Corresponding author: Granville J. Matheson^1^.

## Abstract

PET analysis is conventionally performed as a two-stage process of quantification followed by analysis. We recently introduced SiMBA (Simultaneous Multifactor Bayesian Analysis), a hierarchical model that performs quantification and analysis for all brain regions of all individuals at once, and in so doing improves both the accuracy of parameter estimation as well as inferential efficiency. However until now, SiMBA has only been implemented for the two-tissue compartment model. We have now extended this general approach to also allow a non-invasive reference tissue implementation that includes both the full reference tissue model and the simplified reference tissue model. In simulated data, SiMBA improves quantitative parameter estimation accuracy, reducing error by, on average, 57% for binding potential (*BP*_ND_). In considerations of statistical power, our simulation studies indicate that the efficiency of SiMBA modeling approximately corresponds to improvements that would require doubling the sample size if using conventional methods, with no increase in the false positive rate. We applied the model to PET data measured with [^11^C]AZ10419369, which binds selectively to the serotonin 1B receptor, in datasets collected at three different PET centres (n=139, n=44 and n=39). We show that SiMBA yields replicable inferences by comparing associations between PET parameters and age in the different datasets. Moreover, we show that time activity curve data from different centres can be combined in a single SiMBA model using covariates to control between-centre parameter differences, in order to harmonise data between centres. In summary, we present a novel approach for noninvasive quantification and analysis of PET time activity curve data which improves quantification and inferences, enables effective between-centre data harmonisation, and also yields replicable outcomes. This method has the potential to significantly expand the range of research questions which can be meaningfully tested using conventional sample sizes with PET imaging.

## 1 INTRODUCTION

Quantification of brain Positron Emission Tomography (PET) data involves fitting compartmental pharmacokinetic (PK) models to measurements of radiotracer concentrations in tissue over time, called a time activity curve (TAC). These models describe the transfer of the radiotracer among different binding compartments, or binding states, by estimating the rates of transfer between them. Using these rate constants, we can derive an estimate of the availability of the target molecule, for example binding potential (BP), which describes the specific binding of the radiotracer to the target of interest relative to some reference quantity. The gold standard for PET quantification involves sampling the arterial blood throughout the PET examination to derive a metabolite-corrected arterial input function. This procedure, termed invasive quantification, is labour-intensive, can be uncomfortable for participants, and requires experienced staff for arterial cannulation, blood measurement, as well as analysis. To this end, non-invasive quantification approaches were developed so that specific binding in a target region could be estimated relative to a “reference region”, one that shares properties similar to the region of interest but in which there is no specific binding. Non-invasive quantification therefore tends to be preferred whenever there is a valid reference region and when their estimates can be shown to agree sufficiently with invasive estimates.

In both invasive and non-invasive PET quantification, PK models make a tradeoff between their complexity (i.e. the number of estimated parameters) and their ability to adequately describe the measured data. Simpler models tend to yield parameter estimates which are more stable. However, in order to reduce model complexity, these models must make assumptions about the underlying biology which are not entirely consistent with what is occurring. This can lead to bias in the estimated parameters as well as underfitting the measured data. More complex models are usually able to describe the true underlying PK behaviour of the radiotracer more accurately and may thereby minimise bias in the estimated parameters, however this can also result in reduced stability of parameter estimates. Focusing on non-invasive models, the first developed model was the full reference tissue model (FRTM; Cunningham et al., 1991), which estimates four parameters. This model tends to be quite unstable for most applications, which led to the development of the simplified reference tissue model (SRTM; Lammertsma and Hume, 1996). SRTM includes only three parameters by assuming that equilibrium between specific and non-specific binding compartments is so rapidly established that they can effectively be treated as a single compartment. To this day, SRTM is one of the most widely-used non-invasive models in PET, however even with only three parameters, the model estimates can still be unstable in some applications. This led to the development of SRTM2, which reduces the number of free parameters to two (Wu and Carson, 2002) by assuming that *k*_2′_, the rate at which radioligand leaves the reference tissue to enter the blood, is the same for all target regions of each individual. By simplifying the model in this way, the error in estimates of *R*_1_ and *BP*_ND_ is reduced further, representing the relative delivery rate and binding potential relative to the non-displaceable compartment respectively (Innis et al., 2007). However while this assumption is certainly theoretically true since the reference region is the same for all regions, SRTM2 is nevertheless known to yield more biased outcomes than SRTM (Wu and Carson, 2002; Mandeville et al., 2016).

The conventional approach to using PET to address research questions can be described as a two-stage process: first, binding is quantified in each target region for each individual; and secondly, these binding estimates are entered into a statistical model to draw inferences, for instance comparing patients with healthy controls, or before and after treatment (Chen et al., 2019). While valid, this approach is inefficient in several respects. To address these inefficiencies, we recently introduced SiMBA (Simultaneous Multifactor Bayesian Analysis) (Matheson and Ogden, 2022): a hierarchical, multivariate approach for PET PK modelling which is applied to TAC data. Firstly, SiMBA performs PK quantification of all TACs from all regions of all individuals simultaneously, which allows the model to borrow strength across the sample and exploit similarities in parameter values between individuals or regions. This greatly improves the accuracy of quantification while also reducing the number of *effective* parameters (Gelman et al., 2014; Vehtari et al., 2017), without introducing additional PK assumptions. Secondly, SiMBA performs both quantification and analysis simultaneously as a one-stage model, thereby improving inferential efficiency through more effective error propagation. Lastly, as opposed to conventional practice of retaining only the target binding parameter, SiMBA is able to take advantage of multivariate associations between all of the estimated PK parameters in the statistical model (Matheson and Ogden, 2023). In simulated data, we have shown that this approach not only yields substantial improvements in the accuracy of PK parameter estimates, but also that it greatly improves inferential efficiency with greater statistical power and greater precision of inferences, without increasing the false positive rate (Matheson and Ogden, 2022). However, until now, SiMBA has only been applied to invasive quantification models.

In this study, we introduce a non-invasive implementation of SiMBA, which can be applied either to the SRTM or the FRTM. First, in simulated data, we test its accuracy and inferential efficiency compared to conventional approaches. Next, we evaluate its performance in empirical PET data using [^11^C]AZ10419369, which binds to the serotonin 1B (5-HT^1B^) receptor. This radiotracer was selected for validation of the method because its binding is highly selective, it has a suitable reference region with negligible specific binding, and the validity of reference tissue models for its quantification has been demonstrated (Varnäs et al., 2011). We compare model performance and inferences in data collected at three different PET research centres to assess the consistency of age associations across each centre, as well as the ability of the model to harmonise across TAC data collected at each of the centres in a combined model.

## 2 METHODS

### 2.1 Generalised Model Framework

As previously done for the invasive SiMBA model, we define linear models for each of the log-transformed PK parameters, with partially pooled deviations for each individual, region, and TAC (i.e. the interaction of regions and individuals) which are assumed to be independent from one another (Matheson and Ogden, 2022). By making use of partially-pooled deviations, also called random effects, SiMBA is a hierarchical model (McElreath, 2016; Betancourt, 2020). Moreover, by making use of multiple overlapping hierarchies, SiMBA is a multifactor model (Betancourt, 2021), i.e. regional deviations are estimated across all individuals for each region, while individual deviations are estimated across all regions for each individual. Lastly, by estimating associations between the partially pooled deviations, or random effects, within each of these hierarchies across multiple PK parameters at once, and thereby regularising estimates towards the mean using multivariate shrinkage, SiMBA is a multivariate multifactor model.

This progressive decomposition of the variance of each PK parameter into its separate components has several benefits. Firstly, the majority of variance is explained by parameters estimated over large cross-sections of the data, i.e. across all regions or all individuals, as opposed to estimated independently from each TAC. This has the effect of minimising the “room for error” at the individual TAC level since these deviations are highly regularised, and therefore the SiMBA model is able to give rise to more accurate parameter estimates (Matheson and Ogden, 2022). For instance, in the analysis of [^11^C]WAY100635 data using SiMBA and the two-tissue compartment model, TAC-level deviations accounted for only 2.6% of the total variance in estimated *BP*_ND_ values. The remaining 97.4% of the variance was accounted for by individual and regional deviations which represented only 10% of the total number of parameters [REF SiMBA WAY MDD]. Secondly, by accounting for the majority of the variance using only a small number of parameters in this way, where the large number of deviations at the TAC level are highly-regularised, the number of *effective* parameters of the total model is reduced (Gelman et al., 2014; Vehtari et al., 2017). For instance, when fitting the two-tissue compartment (2TC) model using SiMBA, it was estimated that the total model was fitted using only 2.2 effective parameters per TAC, despite there being 5 parameters estimated for each TAC within the model (Matheson and Ogden, 2022). This has the effect of improving the identifiability and stability of more complex models. Lastly, this approach provides additional explanatory potential as the marginal effects across regions and individuals can be interpreted to understand average effects after compensating for the influence of other factors.

The model estimates each PK parameter (*i*) out of *m* total PK parameters for each individual (*j*), region (*k*) and TAC (*j, k*, i.e. individual × region). The linear model is defined with a global intercept (*α*_*i*_) with unpooled (i.e. fixed effect) covariate deviations 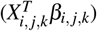 and partially pooled (i.e. random effect) deviations from the expected value for each individual (*τ*_*i, j*_), region (*υ*_*i,k*_) as well as TAC (*ϕ*_*i, j,k*_). By using log-transformed parameters, these deviations represent proportional rather than additive deviations, i.e. an additive deviation of 0.3 from the mean in log *BP*_ND_ for a given individual within the linear model represents an expected value for that individual which is 35% higher than the mean for all regions, before accounting for TAC deviations. For example, if Region A has a mean *BP*_ND_ of 0.1 and Region B has a mean *BP*_ND_ of 3, then after accounting for the individual deviation above, the expected values for *BP*_ND_ will be 0.13 and 4.0 in the two regions respectively. PK parameters (*θ*) are assumed to be drawn from multivariate normal distributions in order to take advantage of the correlation structure between the PK parameters themselves.

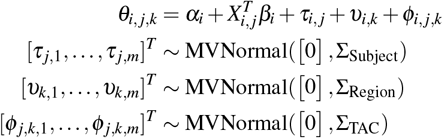

Using these parameters, the data-generating process describing the measured TAC data, *C*_*T*_ (*t*), is defined for each time point (*h*) as follows:

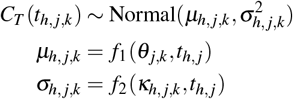

where *f*_1_ represents the PK model itself with which parameters *θ*_*j,k*_, representing the vector of PK parameters for a given individual and region, which are used to generate TAC estimates. Measurement error in the measured TAC data, *σ*, is estimated as a function *f*_2_ of parameters *κ* in the same way as for the invasive SiMBA model (Matheson and Ogden, 2022), i.e. with global mean, partially-pooled deviations across individuals and regions, as well as a series of covariates including the centred natural logarithms of frame duration, region size and injected dose, as well as a non-parametric smooth function over time represented by a thin-plate spline. The former covariates capture the properties of the acquisition and region which influence the measurement error, while the smooth function captures the additional mean pattern of measurement error over the course of the PET examination not accounted for by frame duration, as a result of radiotracer kinetics or other factors. Further details can be found in Matheson and Ogden (2022).

### 2.2 Implementation

#### 2.2.1 Pharmacokinetic Model

For the PK model, represented by *f*_1_, we implemented both the FRTM (Cunningham et al., 1991) and the SRTM (Lammertsma and Hume, 1996)) within the SiMBA framework. The PK parameters of these models can be parameterised in several different ways. In order to facilitate definition of priors, we defined the models to estimate *R*_1_, *k*_2′_ and *BP*_ND_, with the addition of *k*_4_ for the FRTM. We chose to estimate *k*_2′_ as opposed to *k*_2_ because the former is a property of the reference tissue and should, theoretically, be the same, or at least similar, across regions within individuals, and should exhibit less variation between individuals because it should not be influenced by the degree of specific binding. This makes regularisation towards a common mean better justified for this parameter.

When fitting the FRTM to empirical data, the SiMBA model did not converge and yielded *k*_4_ values which appeared to be un-physiological (>1). For this reason, although we implemented the FRTM model, we chose to focus on the SRTM in all following sections.

#### 2.2.2 Modelling of the Reference Tissue

Both FRTM and SRTM involve a convolution with *C*_R_(*t*): the radioactivity concentration in the reference tissue. Numerical convolution tends to be a relatively computationally slow procedure, which is not well-suited for Markov Chain Monte Carlo (MCMC) sampling. For the invasive SiMBA method (Matheson and Ogden, 2022) which involves a convolution of the impulse response function (IRF) with the arterial input function (AIF), we first fit a parametric model to the AIF and then analytically solved the convolution of the IRF with the parametric representation of the AIF to improve computational efficiency. For the non-invasive implementation of SiMBA, we aimed to adopt a similar strategy, however fitting of the reference tissue TAC is not common practice in PET.

To this end, we had to define a model for the reference tissue which was both sufficiently flexible as to faithfully describe the empirical TAC, but also which would make use of functions that lend themselves well to analytical convolution. We opted to define the reference tissue model using the convolution of a Feng parametric AIF model (Wang and Feng, 1992) with a one-tissue compartment model IRF – however in this case, both the parameters of the Feng model and the IRF are estimated using the TAC. For this reason, the parameters of this fit cannot be interpreted biologically – rather, they are only helpful in that they give rise to predicted values which are similar to the reference TAC. This model, which we refer to as the Feng-1TC model, gives rise to 9 parameters: 6 from the Feng model, 2 from the 1TC model, and a *t*_0_ parameter representing the time point before which the AIF is equal to zero. The full working of the Feng-1TC model is described in Supplementary Materials S1. The Feng-1TC model was able to describe the measured reference tissue data well for this (Supplementary Materials S2), and other radiotracers (*not shown*), and has been implemented in the *kinfitr* R package (Matheson, 2019; Tjerkaski et al., 2020). After the development of the Feng-1TC model function, we learned that a similar model has been developed and implemented within the *NiftyPAD* Python software package (Jiao et al., 2023), which only differs in its lacking the *t*_0_ parameter.

The Feng-1TC model was fit to all reference tissue TACs and all fits were visually inspected. Visual inspection is necessary for this model due to the large number of estimated parameters, which can result in occasional poor fits for which clear underfitting is apparent. In all cases, this issue was resolved by randomly selecting new starting parameters from within the upper and lower bounds for the parameters.

#### 2.2.3 SiMBA Model Specification

Section 2.1 laid out the general theoretical framework for the model. However, as for the invasive SiMBA model (Matheson and Ogden, 2022), in practice, this requires several adjustments to account for relevant biological factors. These adjustments were as follows.

The partial pooling of parameters shrinks them towards a common mean. However, due to well-understood anatomical differences in blood flow and regional protein expression patterns, the variation in *R*_1_ and *BP*_ND_ between regions is heterogeneous, and cannot be assumed to follow a conventional statistical distribution in general. For this reason, we make use of unpooled, fixed-effect dummy variables within the covariate matrix to estimate regional differences. This means that these deviations are estimated as independent parameters for each region, as opposed to as being sampled from a shared statistical distribution.

Measurement error, like the PK parameters, is estimated using a global mean and partially pooled deviations from that mean. The magnitude of measurement error in a given TAC is expected to be influenced by factors which are common to each specific PET measurement (e.g. the PET system or the injected dose) or which are common to that particular region (e.g. its relative size), however we do not tend to expect it to be substantially influenced by factors which are specific to a given region within a given PET measurement. For this reason, as well as because regional measurement error is not usually of primary interest in PET studies, we made use of deviations estimated across both individuals and regions for *σ*, but did estimated additional TAC-specific (i.e. individual × region) deviations.

#### 2.2.4 Covariates

Covariates are defined for each of the PK parameters as well as the measurement error. For the within-centre analyses, the covariates were defined as follows. For log *R*_1_, we included only covariates for region (as described in section 2.2.3). For log *k*_2_′, we included covariates for age (centred decades) as an overall proportional change across all regions. For the focal binding parameter, log *BP*_ND_, we defined covariates for region and age (centred decades).

Clinical covariates were defined based on the constitution of the relevant dataset. In datasets that include data from patients, we included a covariate for major-depressive disorder (MDD) patients relative to healthy volunteers (HV), as well as an interaction between age and patient status. When pre-post intervention measurements were performed, we included covariates to account for these within-individual changes as well as changes in symptom scores. Clinical comparisons are not a focus of this paper, but they will be more fully described in an upcoming manuscript (in preparation).

In order to allow for regional variability in the associations with age over and above a global association, we also included partially-pooled region × age deviations from the global covariate, which can also be referred to as random slopes. This allows the model to estimate the average association with these covariates using the fixed effect, but to also account for small differences betweeen regions in the magnitude of the association, for instance to accommodate if binding in some regions declines more rapidly with age than in other regions.

When including data from multiple centres in the same model, we included additional covariates to accommodate centre differences. For all three PK parameters, we included a fixed effect for centre as well as an interaction between centre and region. The former accounts for an overall global mean shift, while the latter allows regions within centres to differ from one another: these differences which could be caused by, for instance, the resolution of the PET system or different preprocessing strategies resulting in slightly different regional constitution.

### 2.3 Model Fitting

The model was implemented using the STAN probabilistic programming language (Carpenter et al., 2017) using brms (Bürkner, 2017). The model code as well as a fully documented analysis notebook is provided within the GitHub repository accompanying this manuscript https://github.com/mathesong/SiMBA_Ref_Materials.

#### 2.3.1 Prior Specification

We aimed to define priors so as not to greatly inform the model, but to exclude domains of parameter space which could be deemed as highly unlikely *a priori*. We defined moderately informative normal priors for the global means in order to ensure that the model initialised in approximately the correct neighbourhood, with standard deviation of 0.25. For deviations across individuals and regions, we used zero-centred regularising priors with standard deviations (SDs) of 0.3 for log *R*_1_ and log *BP*_ND_, and 0.1 for log *k*_2′_. For deviations across TACs, we assigned a SD of 0.025 for all parameters. Fixed effects for centre differences and region × centre interaction effects were assigned a SD of 0.1. Age effects were given a standard deviation of 0.1, while diagnosis and treatment effects were assigned a SD of 0.05. The priors are more fully described in Supplementary Materials S3.

### 2.4 Simulations

In order to generate realistic parameter values, we simulated datasets to resemble the true PET data using the extracted posterior mean values from the model fitted to the largest of the three datasets (from Karolinska Institutet, KI). For variation across individuals and TACS (i.e. region × individual), we sampled parameter values from the relevant univariate and multivariate normal distributions. For variation across regions, we made use of the posterior mean values. In this way, we simulate from the same set of regions, but in a new set of individuals with new variation at the regional level within individuals. The simulation settings are more fully described in Supplementary Materials S4.

Because the reference region TAC must be fitted, but must also be checked for imperfect fits, we generated a reference TAC library. To this end, we first fit all reference TACs within the dataset using the Feng-1TC model and visually inspected the fits. Next, we sampled with replacement from the fits to the subset of 93 minute PET acquisitions to generate *true* reference TACs, to which we added noise to generate a reference TAC library of 500 reference TACs. The noise, or measurement error, in reference TACs was calculated as follows.

Firstly, to add variation in noise across the time course of each reference TAC in a realistic manner, we generated predicted values using the regression coefficients estimated in the SiMBA model applied to the empirical KI dataset for measurement error, *σ*, using the posterior means for frame duration and the smooth function over time. This yields a multiplier for *σ* (or an additive difference from log *σ*) at each frame. Second, to calculate the mean noise for reference TACs, we extracted the residuals from the empirical fits and divided them by the relevant estimated multiplier for frame calculated above. Thirdly, to add variation from PET measurement to PET measurement, we sampled from a normal distribution with a mean of zero and the standard deviation estimated in the SiMBA model for PET-to-PET variability in log *σ*. Fourth, we added all of these deviations in log *σ* together (equivalent to multiplying them together for *σ*) to determine the standard deviation of measurement error for each time point of each reference TAC. Finally, we added noise to each time point of each of the 500 reference TACs by sampling from a normal distribution with a mean of zero and the estimated *σ*. In this way, we generated 500 simulated reference TACs with similar magnitude of noise to the empirical data.

We then fit each of the 500 simulated reference TACs using the Feng-1TC reference model and visually inspected them all to check for poor fits. Hence, for each of the TACs within the simulated reference TAC library, we have i) a set of *true* Feng-1TC parameters, which produce a ii) *true* reference TAC, as well as a iii) simulated *measured* reference TAC and iv) a set of *estimated* Feng-1TC parameters. For simulating TAC data in target regions, we sampled from the reference TAC library, and simulated target TAC data using the *true* Feng-1TC parameters. And when fitting the SiMBA model to the simulated target TAC data, model fitting was performed using the *estimated* Feng-1TC parameters.

TAC data were simulated with equal numbers of HVs and patients, where the patients had a mean log *BP*_ND_ of 0.08 lower than the HVs. We simulated HV TAC data before and after placebo treatment, which did not change the mean log *BP*_ND_ value, although the true regional parameter values still varied on account of the region × individual (TAC) variation. We simulated patient TAC data before and after active treatment, which increases log *BP*_ND_ values by 0.04 before accounting for TAC variation. We evaluated inferential efficiency using the treatment − placebo contrast, i.e. the estimated difference-in-difference value.

## 3 RESULTS

### 3.1 Simulations

#### 3.1.1 Outcome Parameter Estimation

To assess accuracy, we extracted posterior mean estimates of the PK parameters from the baseline measurements of the control group from all of the TAC fits from the simulated data, and calculated metrics of accuracy using the combined data. This amounts to 1190, 2320 and 4320 parameter estimates for each region for SiMBA with *n* = 10, *n* = 20 and *n* = 40 respectively, and 7830 estimates for the NLS comparison. We calculated the root mean squared error (RMSE) of the model estimates compared to the true values as a measure of absolute accuracy, and the Pearson’s correlation with the true values as a measure of relative accuracy. The results are shown in Figure 1.

**Figure 1.**
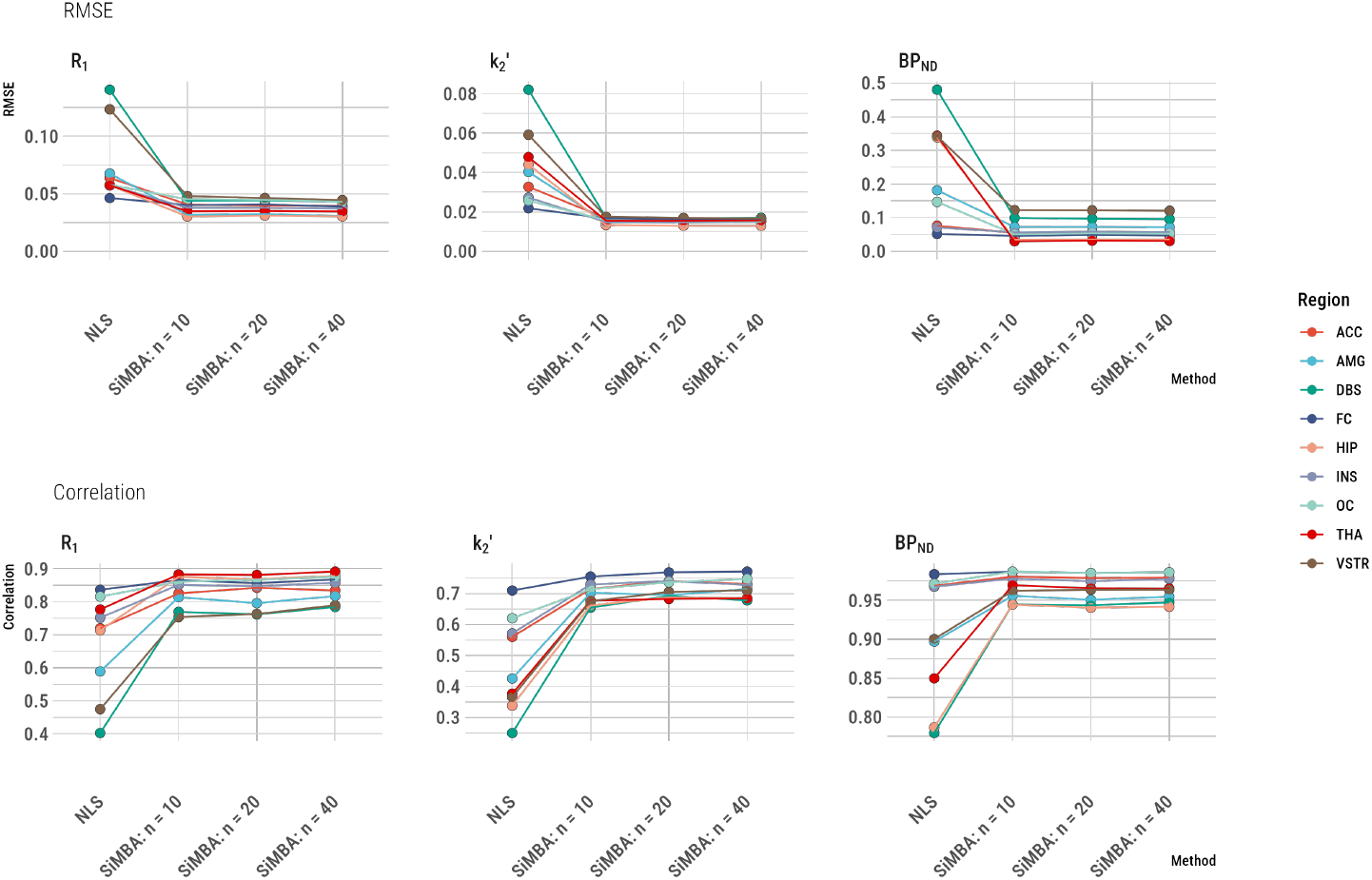
Correspondence between PK parameter estimates and the true values for SiMBA as compared to conventional estimation using nonlinear least squares (NLS). RMSE represents the root mean square error as a measure of absolute deviation from the true values. The correlation values represent the Pearon’s *r* correlation as a measure of the relative accuracy. The number of participants in each of the groups are represented by the *n* values, i.e. *n* = 10 represents 20 participants each measured twice. Regional abbreviations are as follows: ACC is anterior cingulate cortex, AMG is amygdala, DBS is dorsal brain stem, FC is frontal cortex, HIP is hippocampus, INS is insula, OC is occipital cortex, THA is thalamus, and VSTR is ventral striatum.

The RMSE and correlation with the true parameters was improved for every parameter in every region. The greatest improvements in both metrics were expected in the dorsal brain stem (DBS), which is a small midbrain structure with medium-to-low mean [^11^C]AZ10419369 binding. For this region, the RMSE of parameter estimates was reduced by 69% for *R*_1_, by 79% for *k*_2_′, and by 79% for *BP*_ND_ for estimation using SiMBA with *n* = 10 compared to conventional NLS estimation. Across regions, the mean regional reduction of RMSE for SiMBA compared to NLS was 42% for *R*_1_ (13% - 69%, median 40%), 56% for *k*_2_′ (23% - 79%, median 64%), and by 57% for *BP*_ND_ (12% - 91%, median 64%). Similarly, for the DBS, correlations with the true parameter values are increased from 0.40 to 0.77 for *R*_1_, from 0.25 to 0.65 for *k*_2_′, and from 0.78 to 0.94 for *BP*_ND_. The regional outcome RMSE and correlation values are presented in Supplementary Materials S5.

The hierarchical structure of SiMBA implies that quantitative accuracy should improve with larger sample sizes, however this is not evident in Figure 1. We have presented the same results without the NLS outcomes in Supplementary Materials S6, in which it can be observed that there are subtle overall improvements in the RMSE and correlations with the true parameters for *R*_1_ and *k*_2_′, while *BP*_ND_ appears not to be further improved by additional data, potentially suggesting that it has already reached its maximal accuracy with *n* = 10.

#### 3.1.2 Inferential Efficiency

To estimate the degree of improvement to inferential efficiency, we evaluated the ability of SiMBA to estimate the treatment-minus-placebo effects in the simulated datasets: the results are shown in Figure 2. Linear mixed effects (LME) model performance was evaluated in 500 simulated datasets for each condition. Because of the computational burden of the SiMBA model, it was fit to only 50 datasets per condition, and the power and false positive rates were estimated using logspline density functions (Stone et al., 1997) as described in Matheson and Ogden (2022). For this reason, the power and false positive rate of SiMBA inferences are shown with 95% confidence intervals.

**Figure 2.**
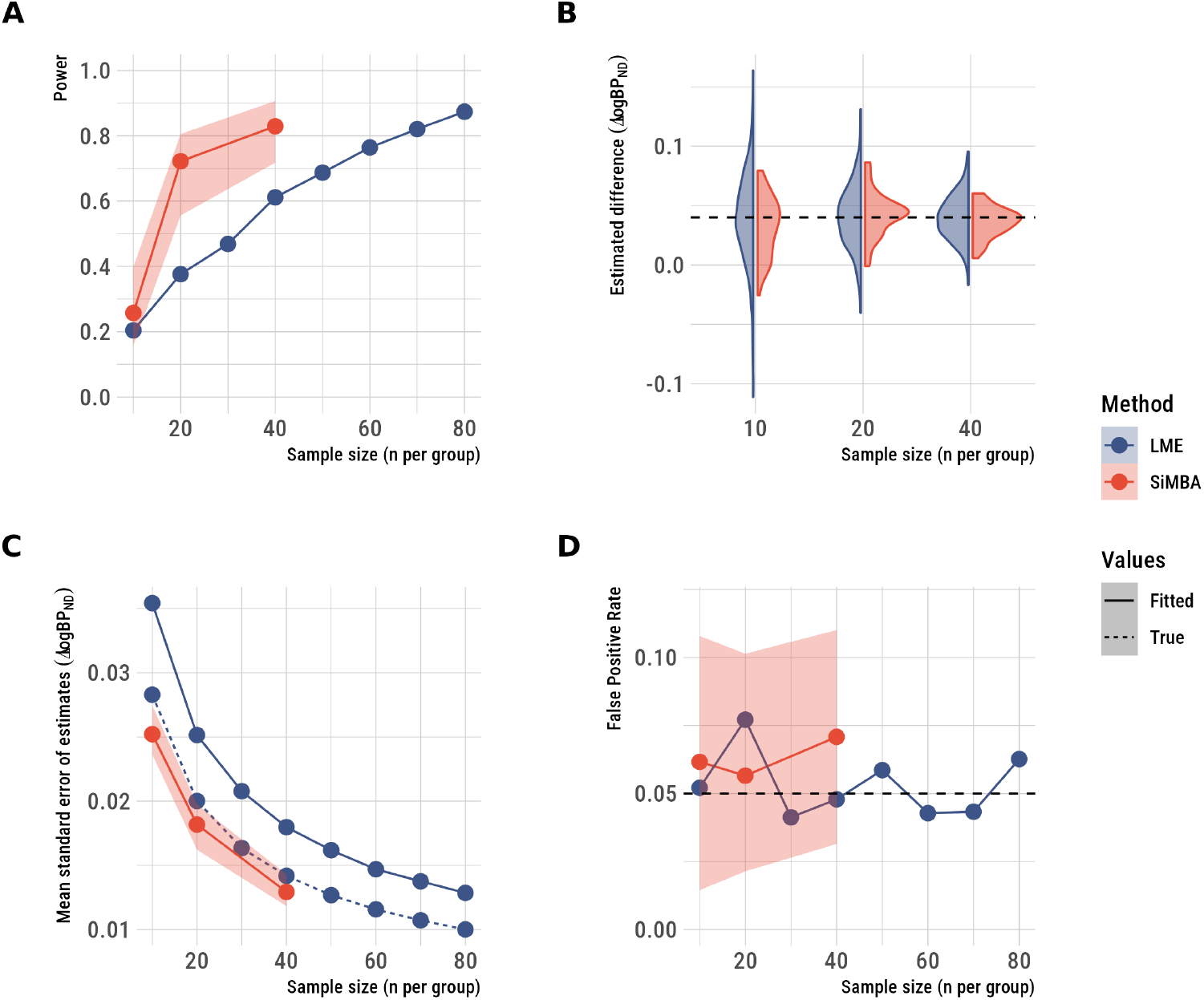
Inferences made using SiMBA as compared to conventional estimation using nonlinear least squares and LME models of log *BP*_ND_ assessing the treatment-minus-placebo contrast. A: statistical power, B. the distribution of the estimated differences across simulated datasets, C the standard error of the estimates, and D. the false positive rate. SiMBA shows improved power, more stable estimates across simulated datasets which are estimated with greater prevision, and no increase in the false-positive rate. The shaded regions represent the 95% confidence intervals.

SiMBA exhibits large improvements to statistical power (Figure 2A), in which conventional methods (using LME) reach the same degree of power with approximately double the sample size for *n* = 20 and *n* = 40. However the improvements are more modest with *n* = 10. SiMBA even showed greater performance than the LME model applied to the true log *BP*_ND_ values, as for the invasive SiMBA model (Matheson and Ogden, 2022). The reason for this is, as we have previously shown Matheson and Ogden (2023), that SiMBA exploits the multivariate relationships between parameters to improve performance, while that information is not considered in a univariate LME analysis applied to the true log *BP*_ND_ values without consideration of the other PK parameters.

The SiMBA model estimates of the treatment-minus-placebo contrast showed less variation across simulated datasets compared to conventional methods, with minimal bias relative to the true value (Figure 2B). Similarly, the standard error of effect estimates is lower using SiMBA compared to using univariate LME models fit to either the true or estimated log *BP*_ND_ parameter values (Figure 2C). At the same time, these improvements to inferential efficiency are not accompanied by increases in the false positive rate (Figure 2D), which remained stable at the nominal 5%.

### 3.2 Application in Empirical Data

#### 3.2.1 Data

The sample used in this study consists of [^11^C]AZ10419369 PET measurements performed at three different PET centres, comprising a total of 222 PET measurements. The Karolinska Institutet (KI) (Nord et al., 2014b,a; Tiger et al., 2020) and Nippon Medical School (NMS) (Tiger et al., 2021) datasets consists of baseline measurements of healthy volunteers (HV) and MDD patients, as well as repeated measurements in a subset of the sample following either treatment or placebo, or no treatment (i.e. a test-retest study). The Neurobiology Research Unit at Rigshospitalet (NRU) consists of HV data obtained through the CIMBI database (Knudsen et al., 2016). The datasets and their properties are described in Table 1 and Figure 3.

**Table 1.** Summary statistics for different datasets.

| Attribute | KI | NMS | NRU |
| --- | --- | --- | --- |
| **Acquisition** |  |  |  |
| PET System | Siemens HRRT | Shimadzu Eminence SET-3000GCT/X | Siemens HRRT |
| Preprocessing | Solena | Solena | PETSurfer |
| Frame Sequence | 93 mins: 38 frames<br>63 mins: 33 frames | 93 mins: 28 frames | 90 mins: 45 frames |
| **Sample** |  |  |  |
| PET Measurements | 139 | 44 | 39 |
| Healthy Volunteers | 62 | 15 | 39 |
| MDD Patients | 40 | 16 | 0 |

**Figure 3.**
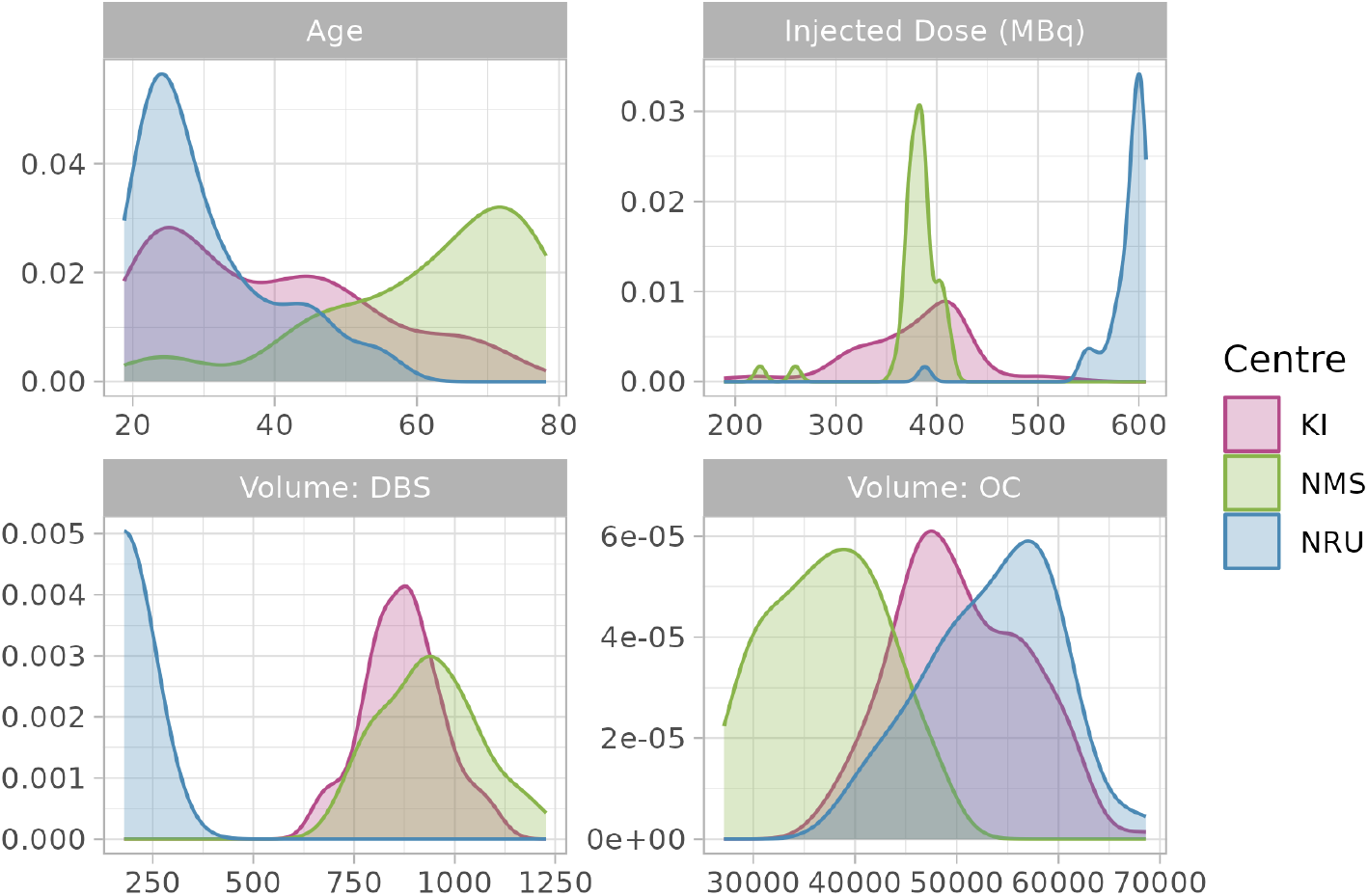
Density plots of age, injected dose of radioactivity and region volumes for different datasets.

Datasets differed from one another in many respects. Most notably, KI and NRU data were acquired using the high-resolution HRRT PET system, while NMS data was collected with the lower-resolution Shimadzu Eminence system. The injected dose was much higher in NRU data as compared to KI and NMS data, and the NMS data measurement frames were longer on average than in the other two datasets. Participant groups at different centres differed a great deal as a function of age: the NRU data was comprised primarily of young individuals, while the NMS dataset was comprised primarily of older individuals. Regarding data preprocessing, KI and NMS data used KI in-house PET analysis software *Solena*, while NRU data were preprocessed using PETsurfer (Greve et al., 2014; Beliveau et al., 2017). These two approaches result in similar region volumes for most regions, however NRU dorsal brain stem (DBS) regions are much smaller as they more specifically delineate the raphe nuclei which lies within the DBS. By including different data collected at different PET centres, we aimed to also assess the ability of our model to harmonise analysis of different data.

#### 3.2.2 Model Fit and Data Harmonisation

The SiMBA-SRTM model was fitted to data collected at each centre independently as well as the combined dataset with covariates to account for centre differences. The model showed excellent fits to the data with highly precise credible and prediction intervals, both within and across datasets (Figure 4). This suggests that the covariates used to accommodate differences in the estimated PK parameters between different datasets with such different properties were sufficient to harmonise the model fit.

**Figure 4.**
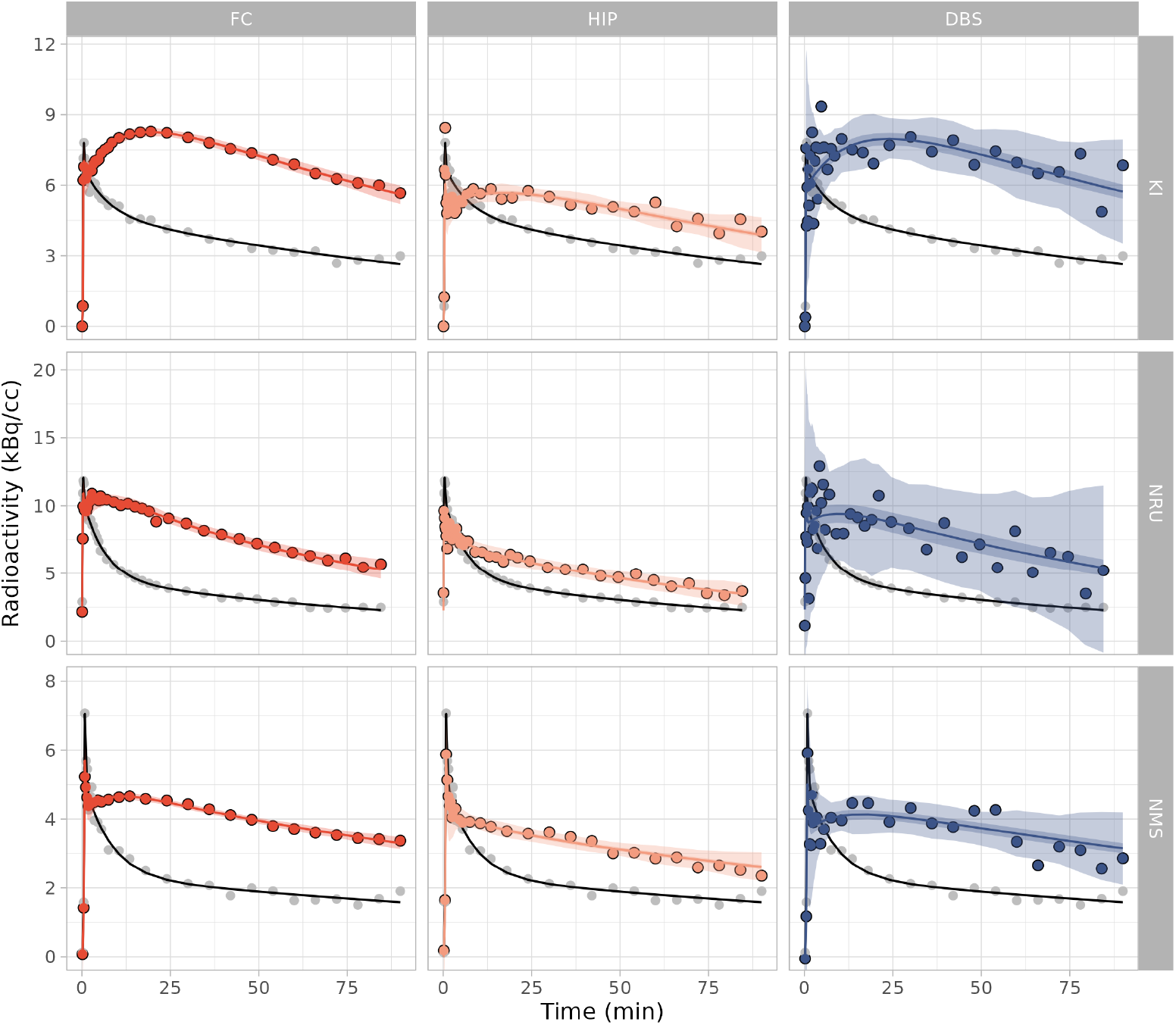
Model fits to the measured data from a randomly selected measurement from each PET centre. These fits are derived from the combined model fit to data from all PET centres, showing fits for the frontal cortex (FC), hippocampus (HIP) and dorsal brain stem (DBS), for data from Karolinska Institutet (KI), the Rigshospitalet (NRU) and Nippon Medical School (NMS). The central line represents the line of best fit, which is surrounded by the shaded region representing the 95% credible interval, which is, in turn, surrounded by the shaded region representing the 95% prediction interval. The grey points represent the measured reference region TAC, with the black line showing the reference region fit which was used for the SiMBA model.

For the combined model which was fit to all data from all centres, the estimated *effective* number of parameters was 4778 for 1984 TACs, equalling 2.4 parameters per TAC. For the largest dataset from KI, in which centre differences did not need to be taken into consideration, the estimated *effective* number of parameters was 2841 for 1243 TACs, equalling 2.3 parameters per TAC. This suggests that while the harmonised model incorporating centre differences required greater model complexity, this additional complexity was minimal.

#### 3.2.3 Consistency of Inferences between Centres: Age

In order to examine the consistency of inferences drawn from its fitting the model to data collected at different research centres, we examined the estimated age associations in data from each centre independently, as well as using the combined model. It has previously been reported not only that [^11^C]AZ10419369 *BP*_ND_ decreases with age, but also that the rate of decrease with age differs between regions (Nord et al., 2014a). For this reason, we compared estimates of both the global mean age association as well as regional variation in the random slopes between regions. We compared model estimates from fitting the model to each centre independently as well as the combined model fit to all the data at once.

As shown in Figure 5, we observe a negative association between *BP*_ND_ and age, as previously reported (Nord et al., 2014a), and in addition, we also find that *k*_2_′ also shows a negative association with age. Regression coefficients representing the proportional rate of change of these parameters per decade shows a high degree of similarity between different centres – despite the difference in age distributions between the NRU and NMS datasets. Furthermore, the combined sample estimate is consistent with all of the individual centre estimates. When comparing the regional differences from the overall mean age effect, the estimates are also highly consistent, from which we conclude that [^11^C]AZ10419369 *BP*_ND_ exhibits more rapid reductions with age in the dorsal brain stem, and less rapid in the ventral striatum and thalamus compared to the mean rate of change. Regional variation in the association between age and *BP*_ND_, assessed by the random slope coefficients, were highly correlated with one another across regions: *r* = 0.79 between KI and NMS, *r* = 0.81 between NMS and NRU and *r* = 0.57 between KI and NRU. The combined model estimates were also highly correlated with the individual research centre estimates: *r* = 0.96 for KI, *r* = 0.92 for NMS and *r* = 0.73 for NRU.

**Figure 5.**
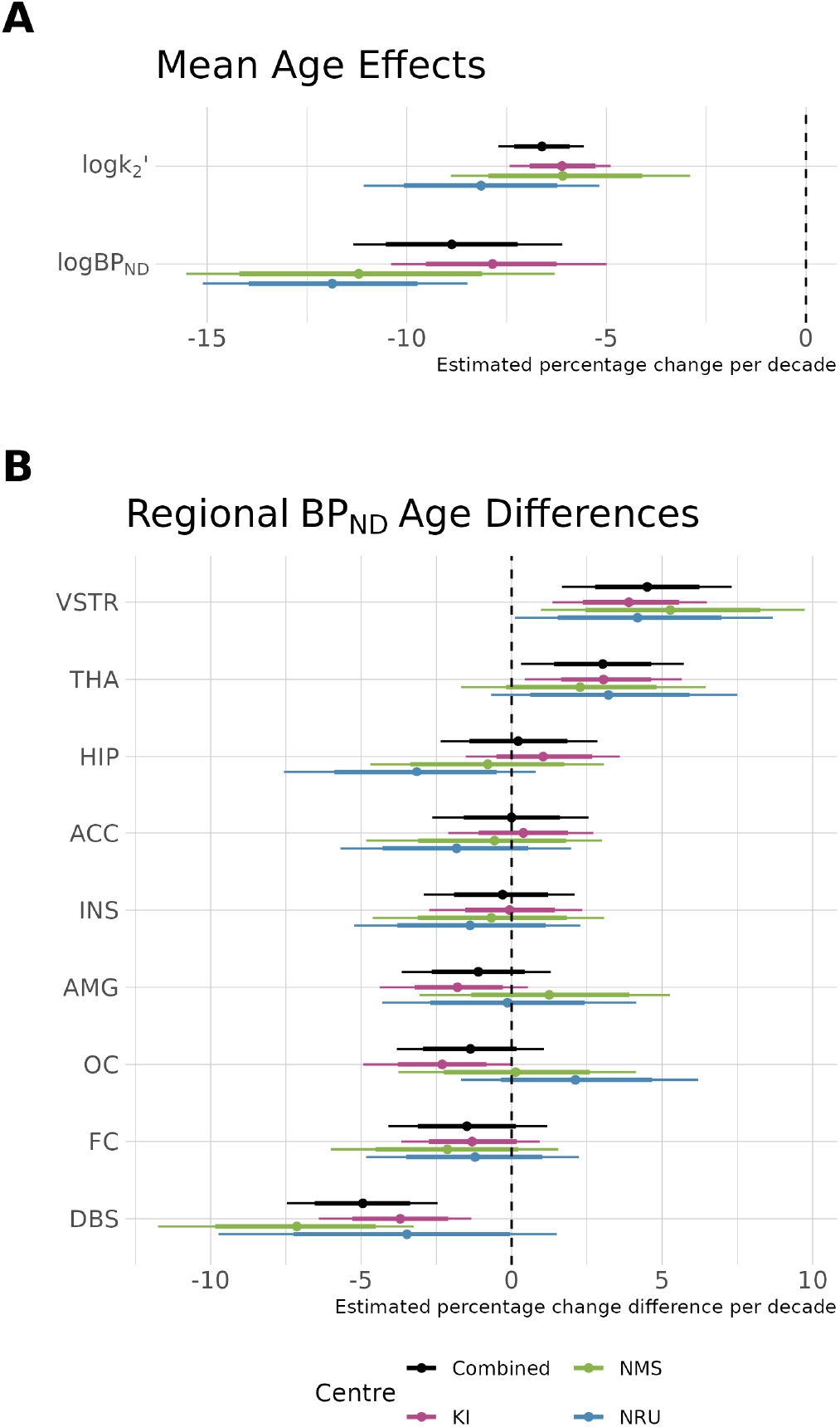
Association of age with PK parameters. A. Overall mean percentage changes in *k*_2_′ and *BP*_ND_ per decade and their estimates in data from each centre independently as well as the combined dataset. B. Regional deviations from the overall mean age effect from the global mean for each region in the decrease in *BP*_ND_ per decade. Regional abbreviations are as follows: ACC is anterior cingulate cortex, AMG is amygdala, DBS is dorsal brain stem, FC is frontal cortex, HIP is hippocampus, INS is insula, OC is occipital cortex, THA is thalamus, and VSTR is ventral striatum. Thick error margins represent the 80% credible intervals while the thinner error margins represent the 95% credible intervals.

#### 3.2.4 Consistency between Centres: k_2_^′^

The PK parameter *k*_2_′ represents the rate of clearance within the reference region. Theoretically, this parameter should be equal for all regions within each individual because the same reference region is used. However, previous studies have shown that setting *k*_2_′ to a common value with the 2-parameter SRTM2 model reduces variance, but also introduces bias relative to the 3-parameter SRTM model in which this parameter is fitted for each region (Wu and Carson, 2002; Mandeville et al., 2016). This suggests that, even though this parameter should theoretically be consistent between regions, the SRTM2 model itself is underfitting the data. We defined the priors for the SiMBA-SRTM model such that, *a priori*, no variation between regions is the most likely outcome, and that greater variation between regions are progressively less likely. To this end, we additionally examined the estimated regional deviations in *k*_2_′, and the degree to which these are consistent between centres.

For all three centres, the model estimated similar degrees of regional variation in *k*_2_′ estimates: the standard deviation of log *k*_2_′ estimates was 0.13 [95% CI 0.08 - 0.20] for NRU, 0.10 [95% CI 0.06 - 0.16] for NMS, 0.12 [95% CI 0.08 - 0.18] for KI, and 0.11 [95% CI 0.07 - 0.17] for the combined sample. In addition, we find that regional *k*_2_′ estimates are also highly consistent between research centres (Supplementary Materials S7), with lower *k*_2_′ in the hippocampus, and higher *k*_2_′ in the frontal cortex and anterior cingulate cortex. Regional difference estimates are highly correlated with one another: *r* = 0.64 for KI and NMS, *r* = 0.53 for NMS and NRU and *r* = 0.75 for KI and NRU; and combined model estimates are highly correlated with the individual research centre estimates: *r* = 0.96 for KI, *r* = 0.79 for NMS and *r* = 0.83 for NRU. This provides further evidence that regional variation in *k*_2_′ is highly replicable and systematic and cannot simply be attributed to noise, and suggests that the SiMBA-SRTM approach is accurately describing and accounting for these differences across different datasets.

#### 3.2.5 Consistency between Centres: Covariance Matrices

The SiMBA framework models and exploits multivariate assocations between the estimated PK parameters to improve performance, and we have previously shown that individual-level covariance matrices across participant groups collected at the same research centre are highly consistent for invasive SiMBA modelling (Matheson et al., 2024). Here we sought to investigate to what extent these matrices are consistent across data collected at different research centres using the same radiotracer. While correlation estimates were somewhat consistent, estimates were less consistent than would be expected based on their credible intervals (Supplementary Materials S8). Between-parameter correlations were most similar between KI and NRU data which were both acquired using HRRT PET systems. Across all model fits, log *BP*_ND_ and log *R*_1_ were highly correlated with one another, as previously reported in (Matheson and Ogden, 2023) for all tested radiotracers, however associations with log *k*_2_′ were less consistent, particularly for the NMS dataset.

## 4 DISCUSSION

In this study, we demonstrate the application of a new approach for fitting PET TAC data using non-invasive PK models with a reference tissue. This approach makes use of hierarchical Bayesian modelling to take advantage of similarities across both regions and individuals to improve estimation accuracy, while also making use of a multivariate framework to exploit between-parameter correlations. By implementing SRTM using this model framework, we show using simulated data that SiMBA can greatly improve parameter estimation, reducing error for some regions by up to 70% for *R*_1_ and 90% for *BP*_ND_. Morever, we show large improvements in the inferential efficiency of these models, resulting in similar statistical power to samples of approximately double the size analysed using conventional methods – all without increasing the false positive rate.

We also demonstrated the application of this method in real measured data collected at three different research centres with different PET systems, frame sequences, age ranges, preprocessing, injected doses and sample sizes. By incorporating centre differences in the model framework, we could fruitfully apply the model to harmonise multi-centre TAC data, resulting in excellent fits to data from all centres. This shows that SiMBA can be used to combine datasets effectively without needing to smooth out or degrade the signal from any one centre to harmonise the data with data collected at other research centres. SiMBA was even able to effectively harmonise data which were preprocessed using different tools, although more consistent preprocessing between centres would likely improve its performance further. This is particularly relevant given the progress made in recent years to enable PET data sharing, with guidelines (Knudsen et al., 2020), standards (Norgaard et al., 2022), an open database (Markiewicz et al., 2021), and new tools for helping users to prepare, process and share their data (Gorgolewski et al., 2017a; Galassi et al., 2024). Next, by examining the estimated association of PK parameters with age, we could show a high degree of consistency in the estimated regression coefficients. In doing so, we show that SiMBA produces scientific conclusions which are replicable and highly consistent between centres in real data. The consistency of these results is particularly compelling given the different age ranges of the three datasets.

Although the multivariate correlation matrices were not as consistent as anticipated between datasets, the consistency of the results examining the relationship between age and *BP*_ND_ and *R*_1_ suggest that this did not greatly affect the generalisability of the conclusions of the model. This does, however, suggest that the multivariate correlation matrices do not solely represent biological relationships between PK properties, but that they may also be influenced by factors which differed between the data sets.

The application of this model to empirical data showed replicable decreases with age of *BP*_ND_ alongside consistent regional heterogeneity. This replicates previous findings using both the same radiotracer (Nord et al., 2014a), as well as using the other main 5-HT_1B_ receptor radiotracer [^11^C]P943 (Matuskey et al., 2012), both of which reported that neocortical regions showed more pronounced age-related decreases than did subcortical or striatal regions. In the present analysis, we extended previous findings to demonstrate both that these decreases are most pronounced in the DBS, and additionally that *k*_2_′ values are also reduced with age, both of which have not previously been reported. The origin of the regional heterogeneity in age-associated decreases is not well understood, although could potentially be explained by the varying proportions of 5-HT_1B_ auto- and heteroreceptors in different regions (Svensson et al., 2022).

Hierarchical models and their strategy of partial pooling can be thought of as a statistically principled method for adaptive regularisation. The degree of regularisation which is required is estimated by the model together with the regularised parameters themselves (McElreath, 2016). In this way, such models effectively balance bias with variance without needing to simplify the biological complexity of the PK models themselves any further, through making the assumption that PK parameters follow defined statistical distributions within the population being examined. The resulting complexity of the model is captured by the effective number of parameters which, for these models was reduced to less than 2.5 parameters per TAC, while also including all covariates, statistical comparisons and modelling the measurement error simultaneously. Hierarchical modelling has been applied to fit TAC data in several previous studies, between individuals (van Rij et al., 2005; Syvänen et al., 2011; Kågedal et al., 2012; Chen et al., 2019; D. Shieh et al., 2024) and between regions (Zamuner et al., 2002; Berges et al., 2013; Kågedal et al., 2012), however to our knowledge, SiMBA is the only approach with which PET TAC data can be modelled simultaneously across both individuals and regions, and in a manner which can be applied to effectively any study design (provided there are sufficiently many participants).

Based on the ability of hierarchical models to simplify estimation of complex PK models, we anticipated that SiMBA would also stabilise estimation of the FRTM (Cunningham et al., 1991) – however the estimated *k*_4_ values were much higher than anticipated (>1), which was a cause for concern, and so we did not proceed with this model. Application of the FRTM may be more successful for other radiotracers for which SRTM is unable to sufficiently describe the data, or else may be improved through setting more informative priors. In any case, for modelling [^11^C]AZ10419369 data, the SRTM appears to describe the data sufficiently well.

The primary disadvantages of the SiMBA model framework is its computational burden. The combined model fitted to all 222 measurements, with 1000 MCMC iterations per chain which is reasonably low, required approximately 6.5 days of sampling on three cores (sampling time scales approximately linearly with sample size and MCMC iterations). Although this is a long time relative to existing models, it is trivial in comparison to the months to years it takes to acquire such large datasets. Another issue is that the implementation of SiMBA requires an understanding of the R programming language and the ability to define reasonable priors. To this end, we have provided fully documented code in an online repository along with an example simulated dataset to assist new users with implementing this model.

In summary, we have introduced a non-invasive implementation of SiMBA that substantially improves both the accuracy of PET parameter estimation and the efficiency of statistical inference. We have further validated its performance, showing that it can yield replicable and precise inferences in independently collected datasets, even when they differ in key properties, and even that it can successfully harmonise data collected at different research centres. For these reasons, we believe that SiMBA is an important addition to the PET modeller’s arsenal as it can expand the range of research questions which can be meaningfully investigated using PET, particularly in cases where sample sizes are limited by practical constraints. Importantly, SiMBA additionally provides a way for existing data to be re-purposed for improving inferences and reducing costs and needless exposure of additional participants to radioactivity. We hope to build a BIDS app (Gorgolewski et al., 2017b) in future to facilitate data preparation and model specification for PET-BIDS datasets (Gorgolewski et al., 2016; Norgaard et al., 2022).

## 5 DATA AND CODE AVAILABILITY

The R code used to apply this method are provided in an open repository (https://github.com/mathesong/SiMBA_Ref_Materials), including a simulated dataset and an annotated notebook demonstrating the method applied to the data. The measured data used in the application section is drawn from previously published studies (Nord et al., 2014a,b; Knudsen et al., 2016; Tiger et al., 2020, 2021).

## 6 ETHICS

All subjects gave written informed consent prior to participation, and all studies were approved by the regional ethics committees.

## 7 AUTHOR CONTRIBUTIONS

**Granville J. Matheson:** Conceptualization, Methodology, Software, Validation, Formal analysis, Investigation, Resources, Data Curation, Writing - Original Draft, Writing - Review & Editing, Visualization, Funding acquisition. **Johan Lundberg:** Conceptualization, Writing - Review & Editing. **Martin Gärde:** Investigation, Data Curation, Writing - Review & Editing. **Emma R Veldman:** Investigation, Data Curation, Writing - Review & Editing. **Amane Tateno:** Writing - Review & Editing. **Yoshiro Okubo:** Writing - Review & Editing. **Mikael Tiger:** Conceptualization, Investigation, Resources, Data Curation, Writing - Original Draft, Writing - Review & Editing, Supervision, Project administration, Funding acquisition. **R. Todd Ogden:** Conceptualization, Methodology, Investigation, Resources, Writing - Original Draft, Writing - Review & Editing, Supervision, Project administration, Funding acquisition.

## 10 ACKNOWLEDGEMENTS

We would like to acknowledge the CIMBI Database for sharing the NRU dataset with us, and particularly Peter Steen Jensen and Vincent Beliveau. We would also like to thank members of the MIND research group at NYSPI and the PET group at Karolinska Institutet for their helpful comments and feedback on the results.

## 11 SUPPLEMENTARY MATERIALS

### 11.1 Supplementary Materials S1: Model Definitions and Analytical Solutions

The Feng model for the AIF is defined as follows:

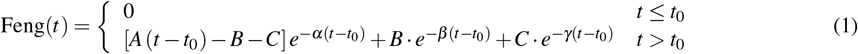

with free parameters *A, B, C, α, β, γ* and *t*_0_.

To derive the estimated reference tissue TAC, *C*_*R*_(*t*), the hypothetical AIF described by the Feng model, Feng(*t*), is convolved with a hypothetical 1TC IRF, IRF

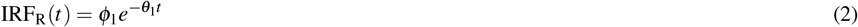

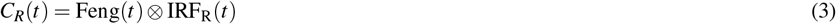

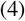

where the 1TC IRF free parameters are *ϕ* _1_ and *θ* _1_ following the terminology of Gunn et al. (2001). The analytical solution of this convolution is as follows:

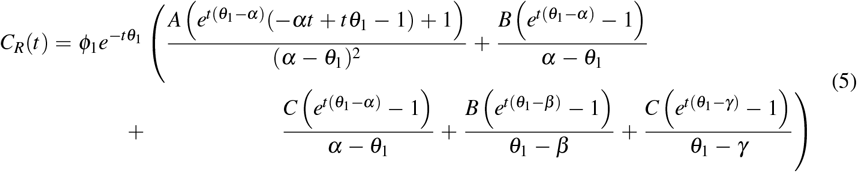

This function is fit to all reference tissue TACs to define a parametric representation of these curves which can be entered into the PK model.

For both the FRTM and SRTM PK models, the convolution within the model is of *C*_*R*_(*t*) with an exponential decay function whose decay is a property of the other parameters of the model, here defined as *c, d* and *q*.

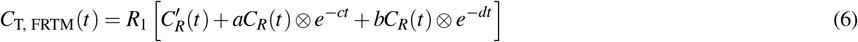

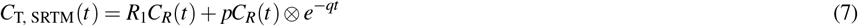

In order to create a general analytical solution of the reference tissue model, we solved the convolution between the reference tissue model with a general exponential decay function which we call ED(*t*) with rate *λ*.

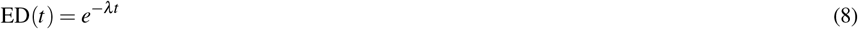

The solution to the convolution of these two functions is then described as follows:

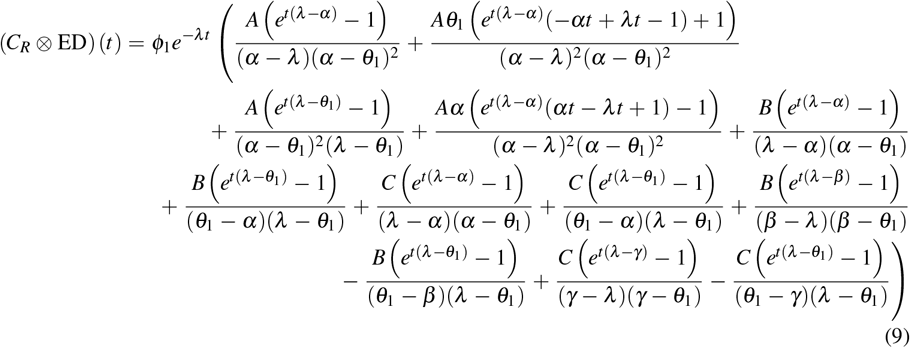

### 11.2 Supplementary Materials S2: Examples of Reference Tissue Fits

**Figure S1.**
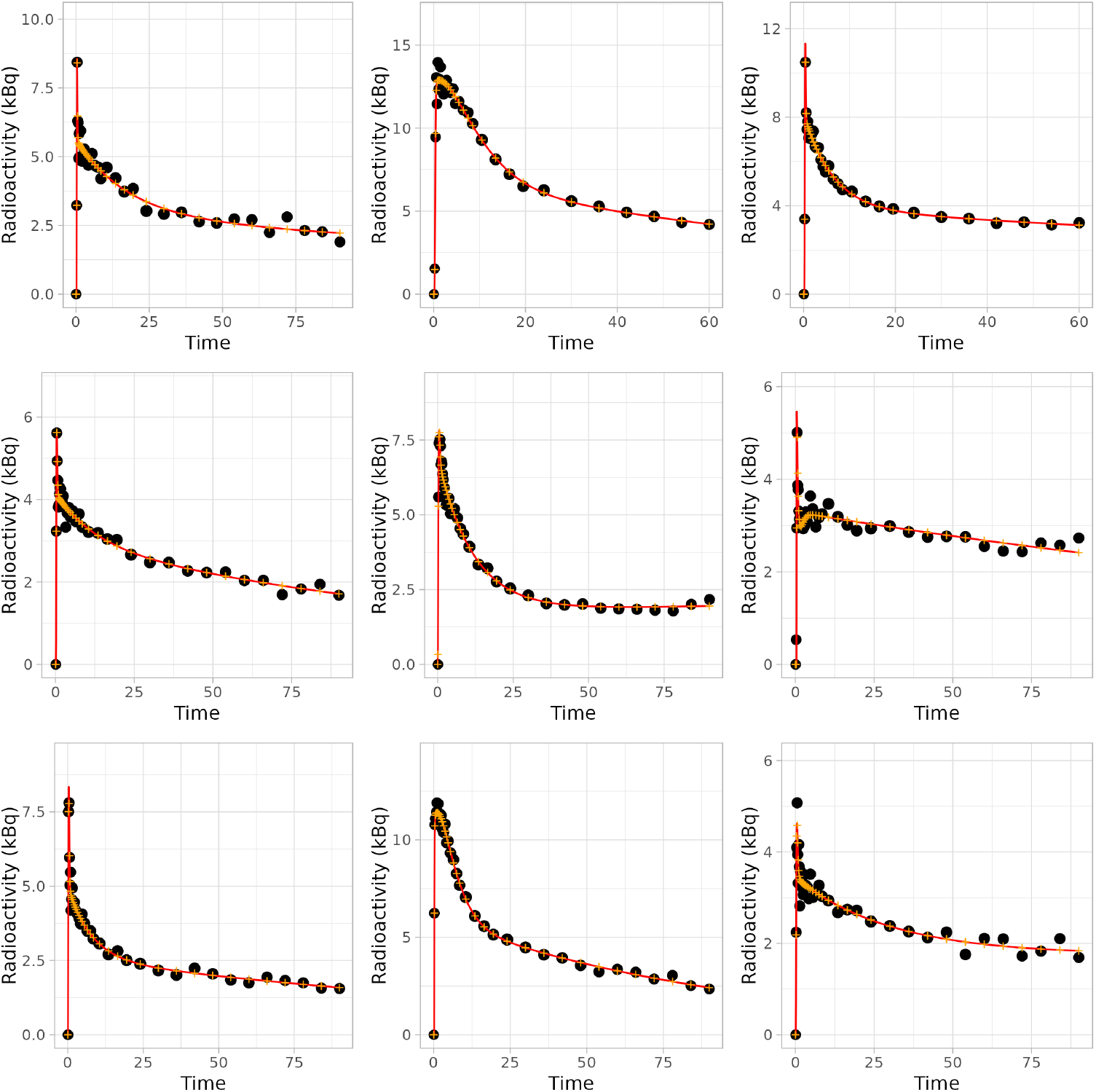
Examples of fits to the cerebellar reference region time activity curves from nine randomly sampled measurements. The size of the data points represents their assigned model weights, and the yellow crosses represent the instantaneous estimated *C*^REF^(*t*) at the mid-frame time points which are used in the PK model functions.

### 11.3 Supplementary Materials S3: Prior Definitions

#### 11.3.1 Global Intercepts

Below are the priors defined for the global intercepts. Note that all priors are defined over the natural logarithms of the parameters.

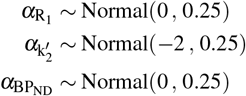

The priors for 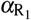 and 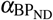 are defined for the frontal cortex as the reference level of the dummy variable.

#### 11.3.2 Individual deviations

Differences between individuals were defined by specifying the primary pharmacokinetic parameters in one variance-covariance matrix.

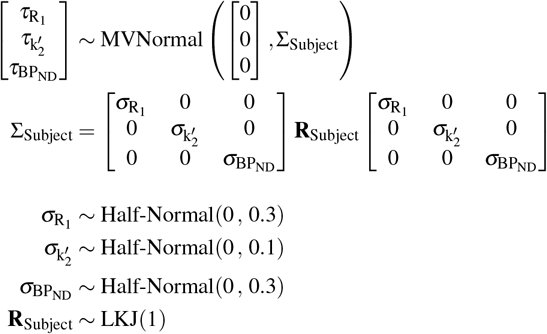

#### 11.3.3 Regional deviations

For log *BP*_ND_ and log *K*_1_, regional differences were defined as unpooled effects using a dummy (indicator) variable defined with reference to the dorsolateral prefrontal cortex. For simplicity, all regional differences (with the exception of [^11^C]DASB) were defined as zero-centred regularising priors with the same SD.

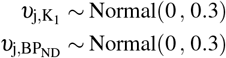

For 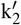, regional differences were defined as pooled variables, arising from a common distribution

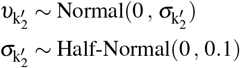

#### 11.3.4 TAC deviations

For the Individual × Region deviations, we made use of highly-constrained deviations

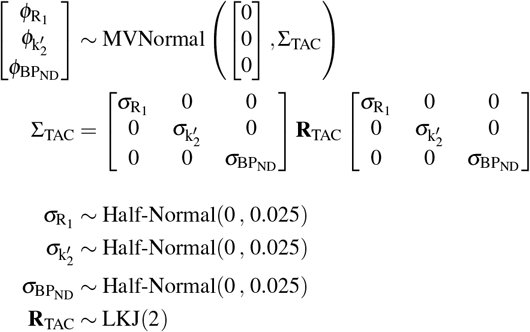

#### 11.3.5 Covariates

Covariate effects were all estimated using zero-centred regularising priors. For the assessment of age, we defined the following priors, for centred age scaled so that a unit change represents a decade.

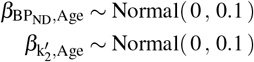

Clinical covariates were defined with wider priors

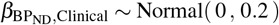

using the same prior for MDD-Control and Treatment-Baseline (ECT, ketamine and placebo) contrasts, as well as for the centred change in symptom scores scaled to a ΔHAM-D of 10 points.

The random variation in slopes between regions was defined using the mean estimate and random slopes derived from a common distribution using the following priors for age:

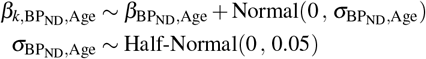

and for clinical covariates:

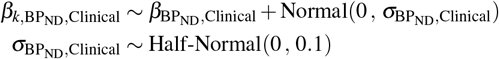

To account for differences between centres, we made use of priors for overall parameter mean shifts, with one parameter estimated for differences to each other centre (i.e. using KI as the reference centre, we estimated a deviation each for both the NRU and NMS datasets).

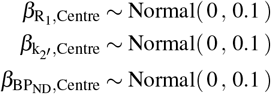

For R_1_ and BP_ND_, we also defined Region × Centre interaction effects to account for differences at the region-within-centre level, with the following priors for each region and centre.

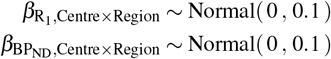

In R code using the *brms* package, the code for defining the model equation and priors is as follows:

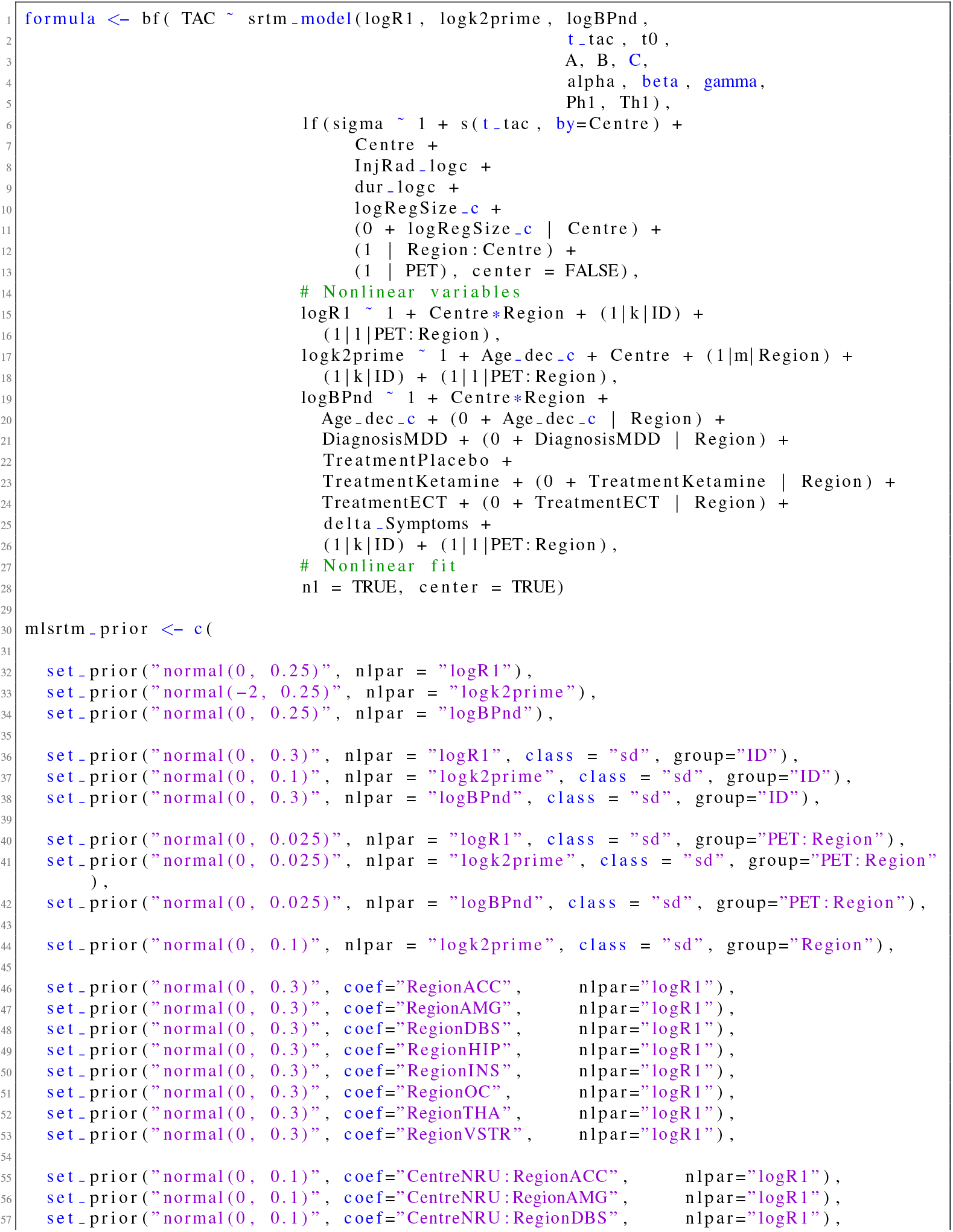
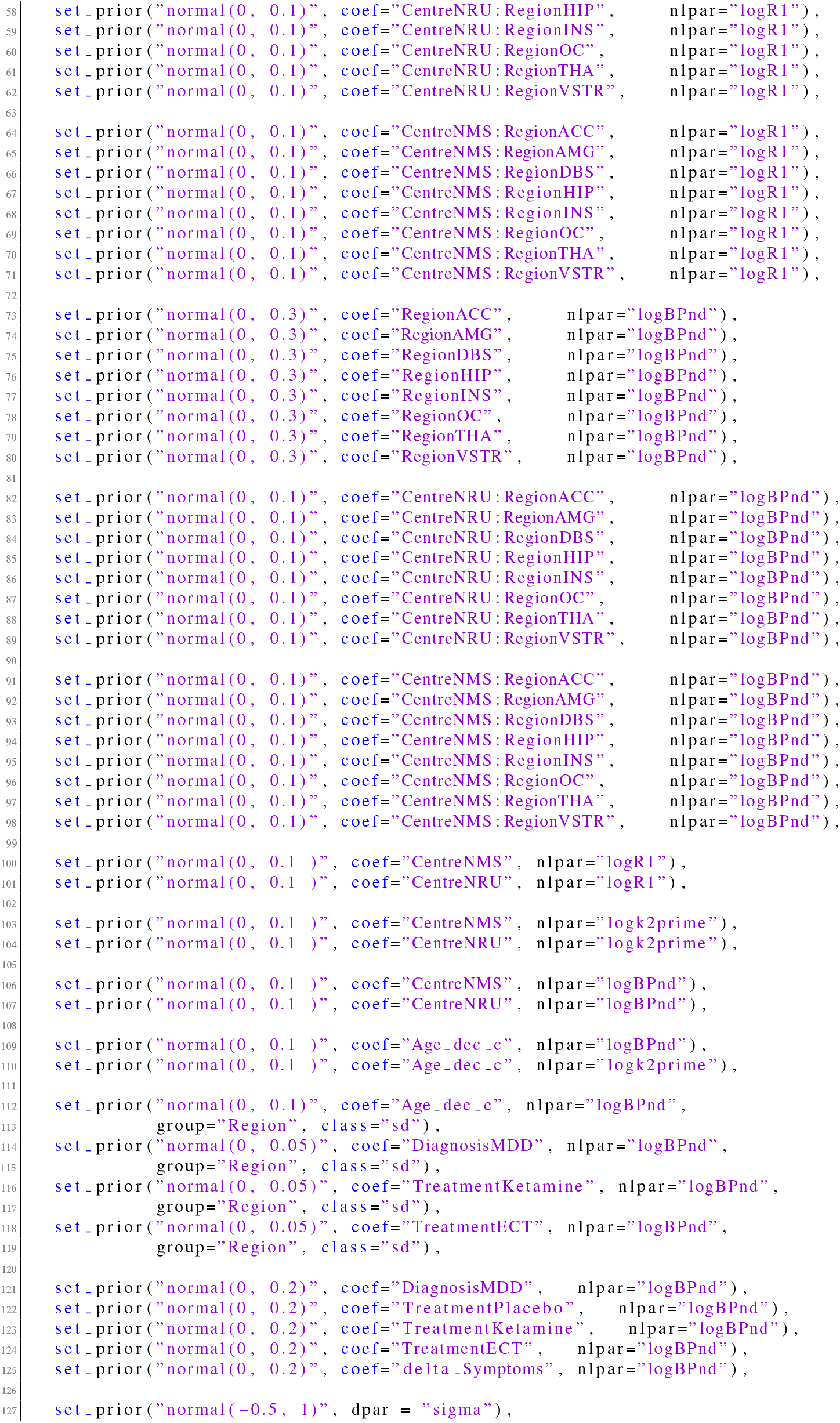
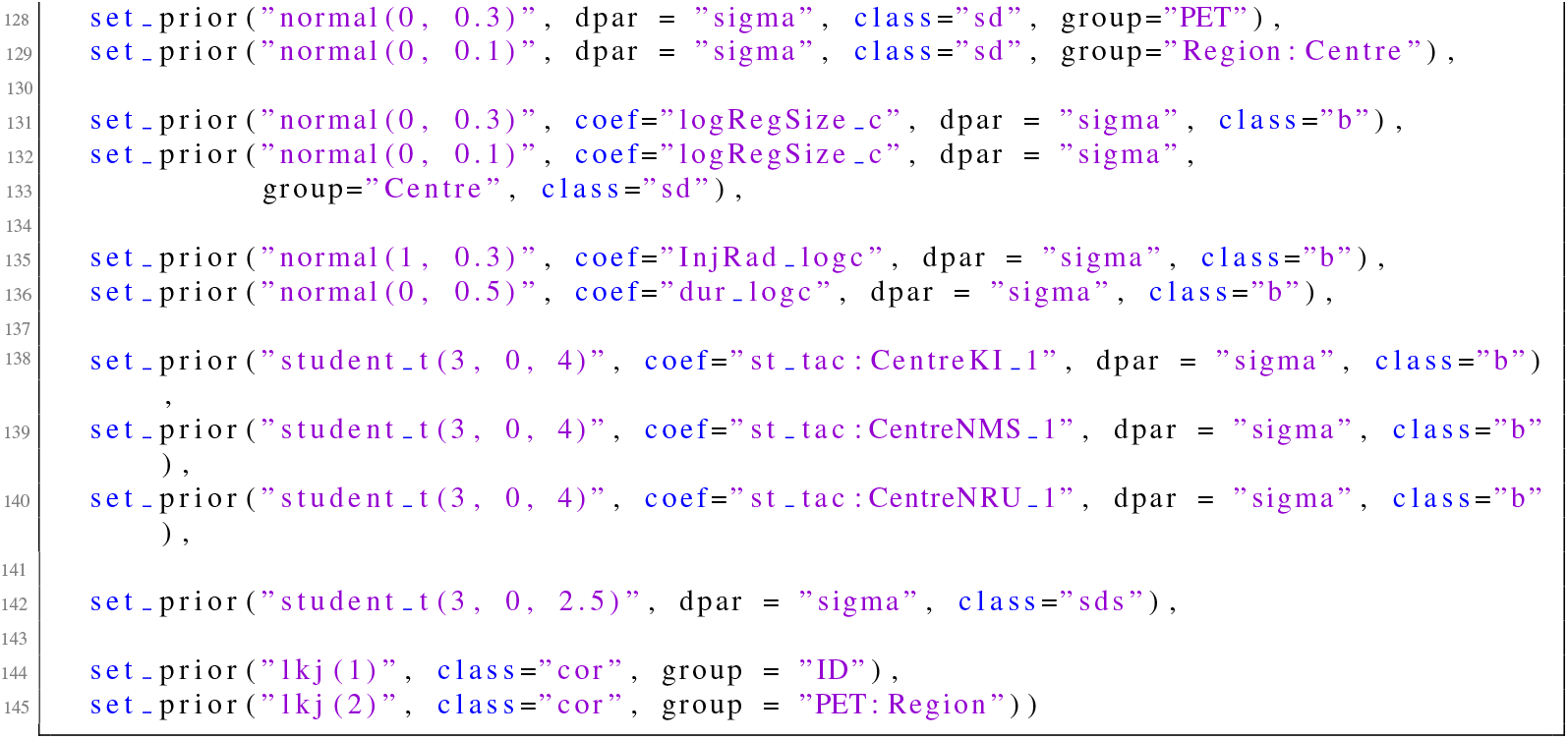

### 11.4 Supplementary Materials S4: Simulation Parameters

#### 11.4.1 Global Intercepts

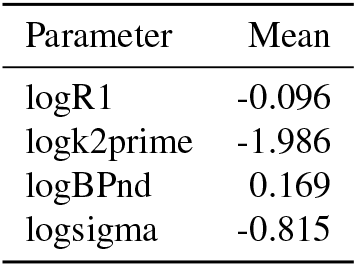

#### 11.4.2 Individual Deviations

These deviations represent the mean individual deviations. When there are two PET measurements within a single individual, there is only a single deviation from the mean defined for that specific individual.

Standard deviation

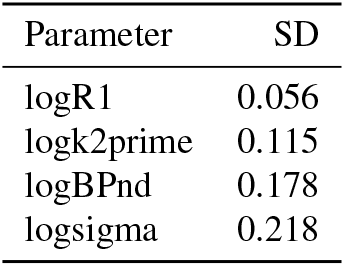

Correlation matrix

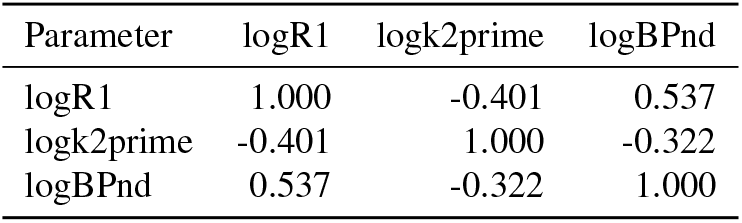

#### 11.4.3 Regional Deviations

For regional deviations, we did not sample from distributions, but rather used the posterior mean deviations for each of the parameters. If we were to sample from distributions instead, we would effectively be simulating a unique set of regions.

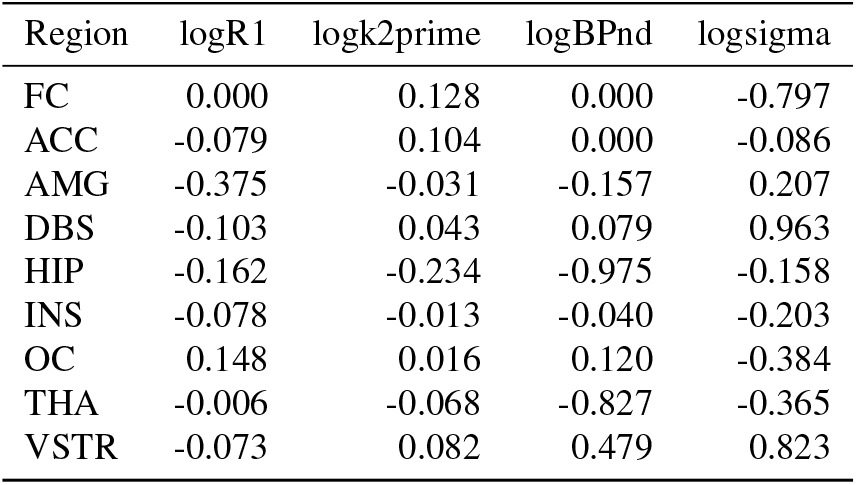

#### 11.4.4 PET x Region Deviations

These deviations were defined for the interaction of the PET measurement and the region. For this reason, they accommodate both PET-to-PET variability as well as Region-within-Individual variability.

Standard deviation

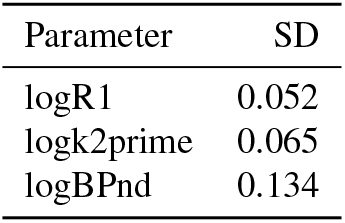

Correlation matrix

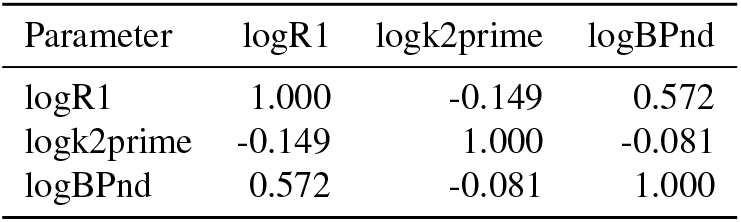

#### 11.4.5 Measurement Error Function

Variation in measurement error, log *σ*, over the duration of the time activity curve was defined with a smooth function and covariates.

The centred smooth function is as follows:

Following addition of measurement error as defined the above function and the region, we also added additional measurement error to account for frame durations by multiplying the centred natural logarithm of the frame duration with the estimated coefficient from the KI dataset of −0.233.

### 11.5 Supplementary Materials S5: Regional improvements in RMSE of PK Parameters

**Figure S2.**
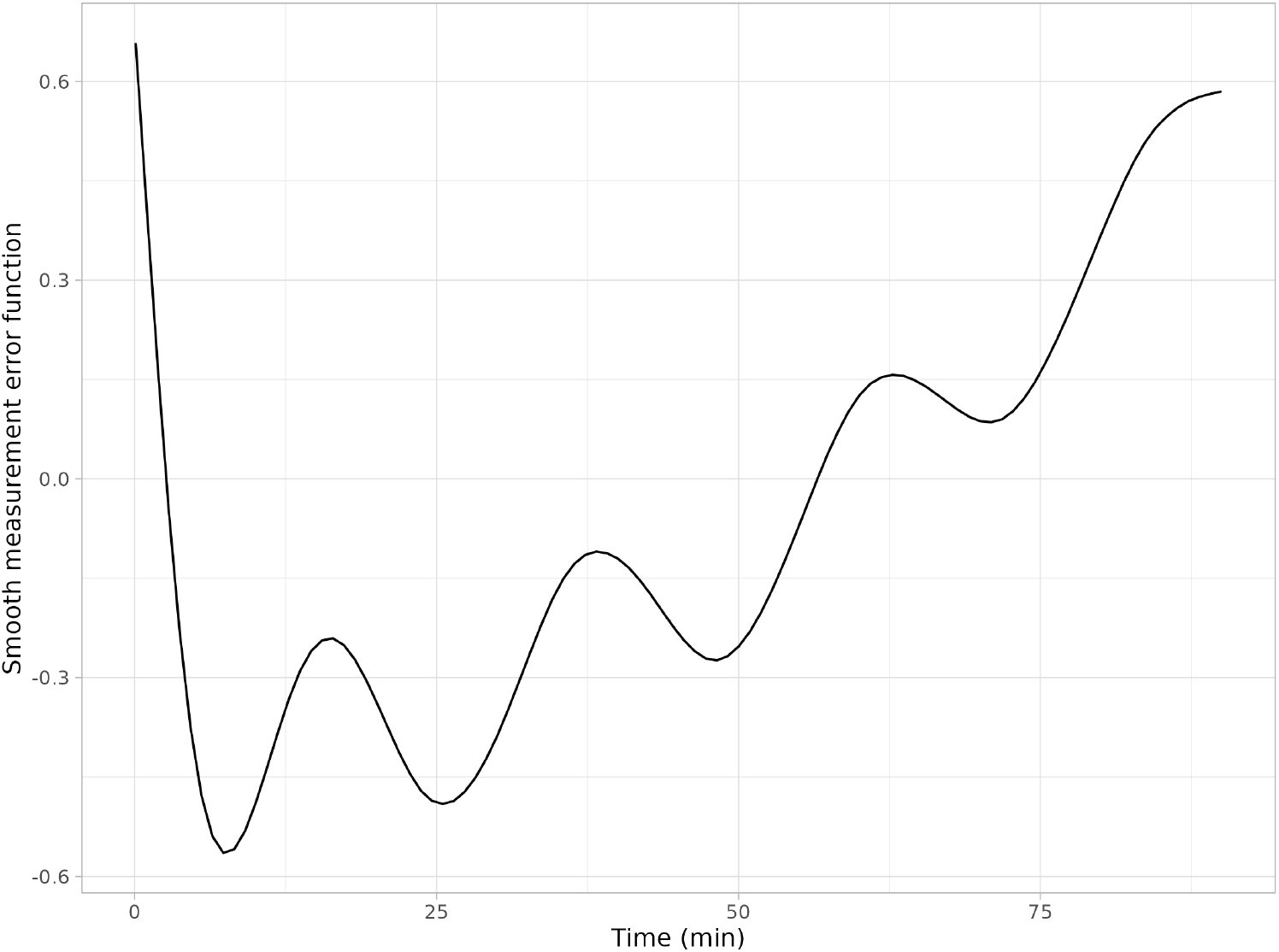
The smooth deviations in the measurement error as a function of TAC time, before accounting for frame duration or region size.

The following table compares the improvements in the RMSE and correlations with the true values for estimation of the following parameters, comparing NLS with SiMBA with n = 10.

RMSE

| Region | NLS | SiMBA | Reduction (%) |
| --- | --- | --- | --- |
| $R_1$ | | | |
| FC | 0.05 | 0.04 | 13 |
| OC | 0.06 | 0.05 | 22 |
| INS | 0.06 | 0.04 | 34 |
| ACC | 0.06 | 0.04 | 36 |
| THA | 0.06 | 0.03 | 40 |
| HIP | 0.06 | 0.03 | 47 |
| AMG | 0.07 | 0.03 | 53 |
| VSTR | 0.12 | 0.05 | 61 |
| DBS | 0.14 | 0.04 | 69 |
| $k_2'$ | | | |
| FC | 0.02 | 0.02 | 23 |
| OC | 0.03 | 0.02 | 39 |
| INS | 0.03 | 0.01 | 46 |
| ACC | 0.03 | 0.02 | 48 |
| AMG | 0.04 | 0.01 | 64 |
| THA | 0.05 | 0.02 | 68 |
| HIP | 0.04 | 0.01 | 70 |
| VSTR | 0.06 | 0.02 | 70 |
| DBS | 0.08 | 0.02 | 79 |
| $BP_{ND}$ | | | |
| FC | 0.05 | 0.05 | 12 |
| INS | 0.07 | 0.06 | 21 |
| ACC | 0.08 | 0.05 | 28 |
| AMG | 0.18 | 0.07 | 60 |
| VSTR | 0.34 | 0.12 | 64 |
| OC | 0.15 | 0.05 | 66 |
| DBS | 0.48 | 0.10 | 79 |
| HIP | 0.34 | 0.03 | 90 |
| THA | 0.34 | 0.03 | 91 |

Correlation

| Region | NLS $R_1$ | SiMBA $R_1$ | NLS $k_2'$ | SiMBA $k_2'$ | NLS $BP_{ND}$ | SiMBA $BP_{ND}$ |
| --- | --- | --- | --- | --- | --- | --- |
| ACC | 0.72 | 0.82 | 0.56 | 0.71 | 0.97 | 0.98 |
| AMG | 0.59 | 0.81 | 0.43 | 0.70 | 0.90 | 0.96 |
| DBS | 0.40 | 0.77 | 0.25 | 0.65 | 0.78 | 0.94 |
| FC | 0.84 | 0.87 | 0.71 | 0.75 | 0.98 | 0.99 |
| HIP | 0.71 | 0.88 | 0.34 | 0.66 | 0.79 | 0.95 |
| INS | 0.75 | 0.85 | 0.57 | 0.73 | 0.97 | 0.98 |
| OC | 0.82 | 0.86 | 0.62 | 0.71 | 0.97 | 0.99 |
| THA | 0.78 | 0.88 | 0.38 | 0.68 | 0.85 | 0.97 |
| VSTR | 0.47 | 0.75 | 0.37 | 0.67 | 0.90 | 0.96 |

### 11.6 Supplementary Materials S6: Parameter Estimation Accuracy for SiMBA estimates only

**Figure S3.**
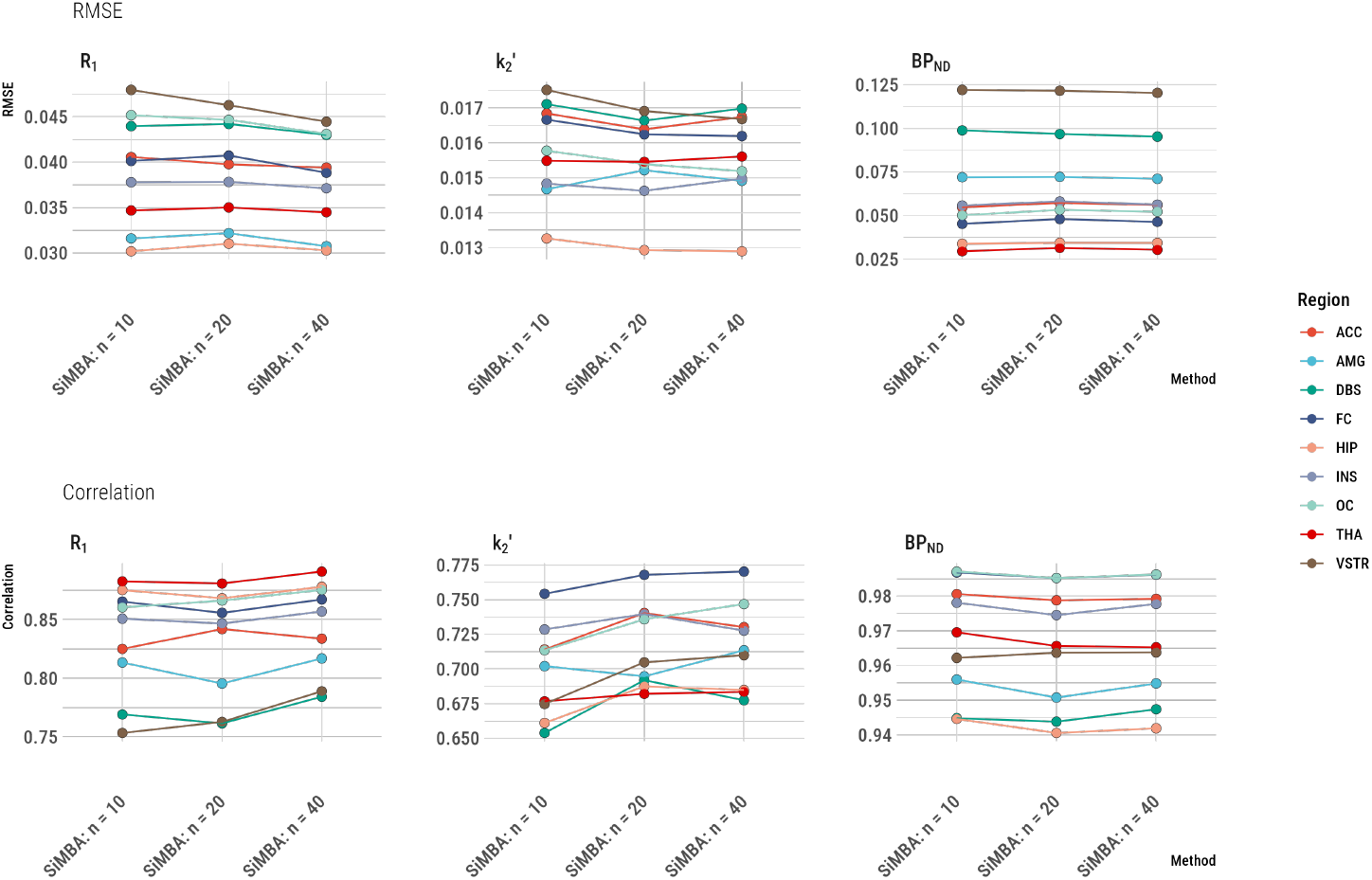
Parameter estimation accuracy assessed by the RMSE and correlation with the true values for SiMBA estimates.

### 11.7 Supplementary Materials S7: Regional *k*_2_′ Estimates

**Figure S4.**
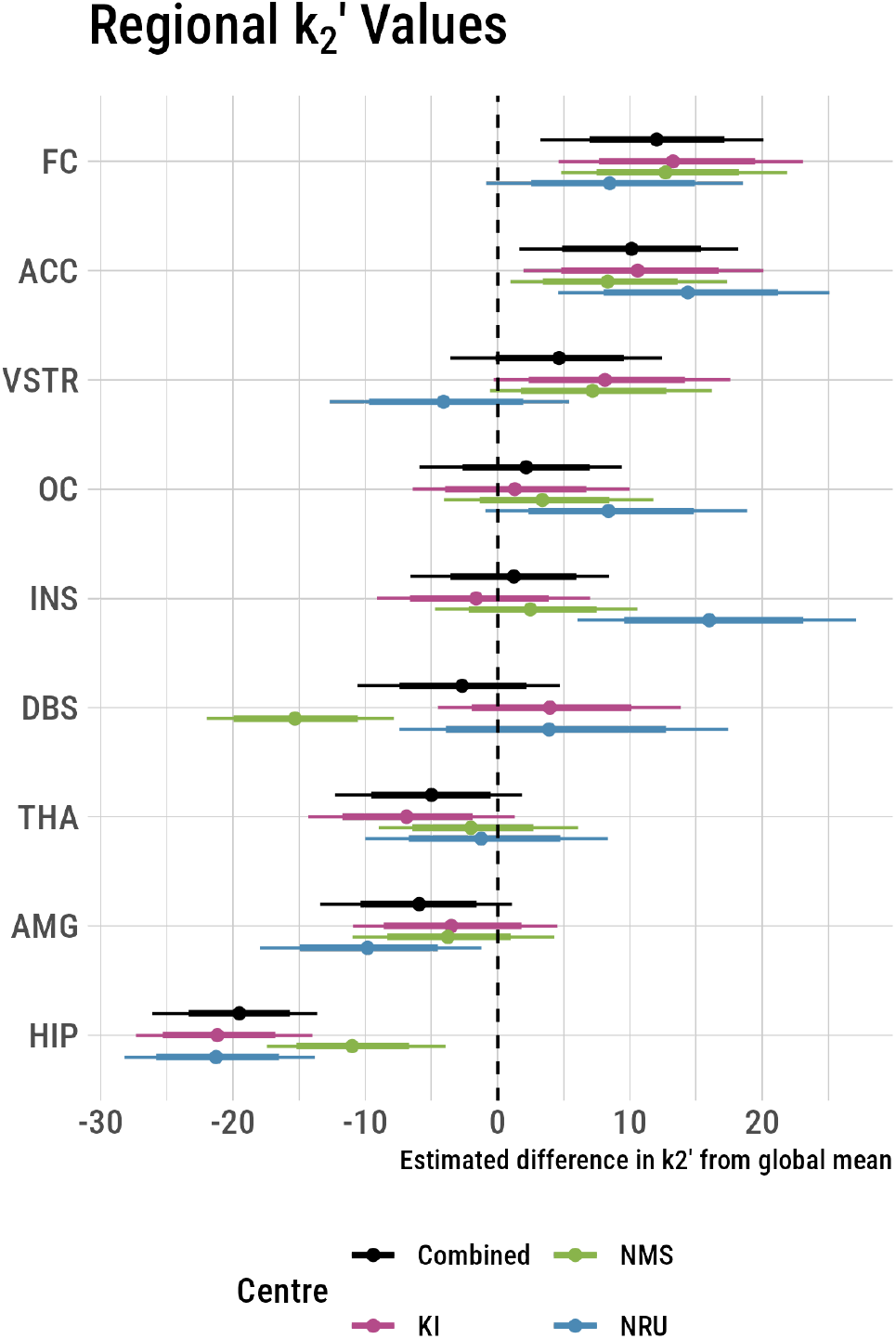
Regional deviations in 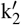 are similar between centres.

### 11.8 Supplementary Materials S8: Correlation Matrices and their Credible Intervals

The individual deviation correlation matrices are shown below.

**Figure S5.**
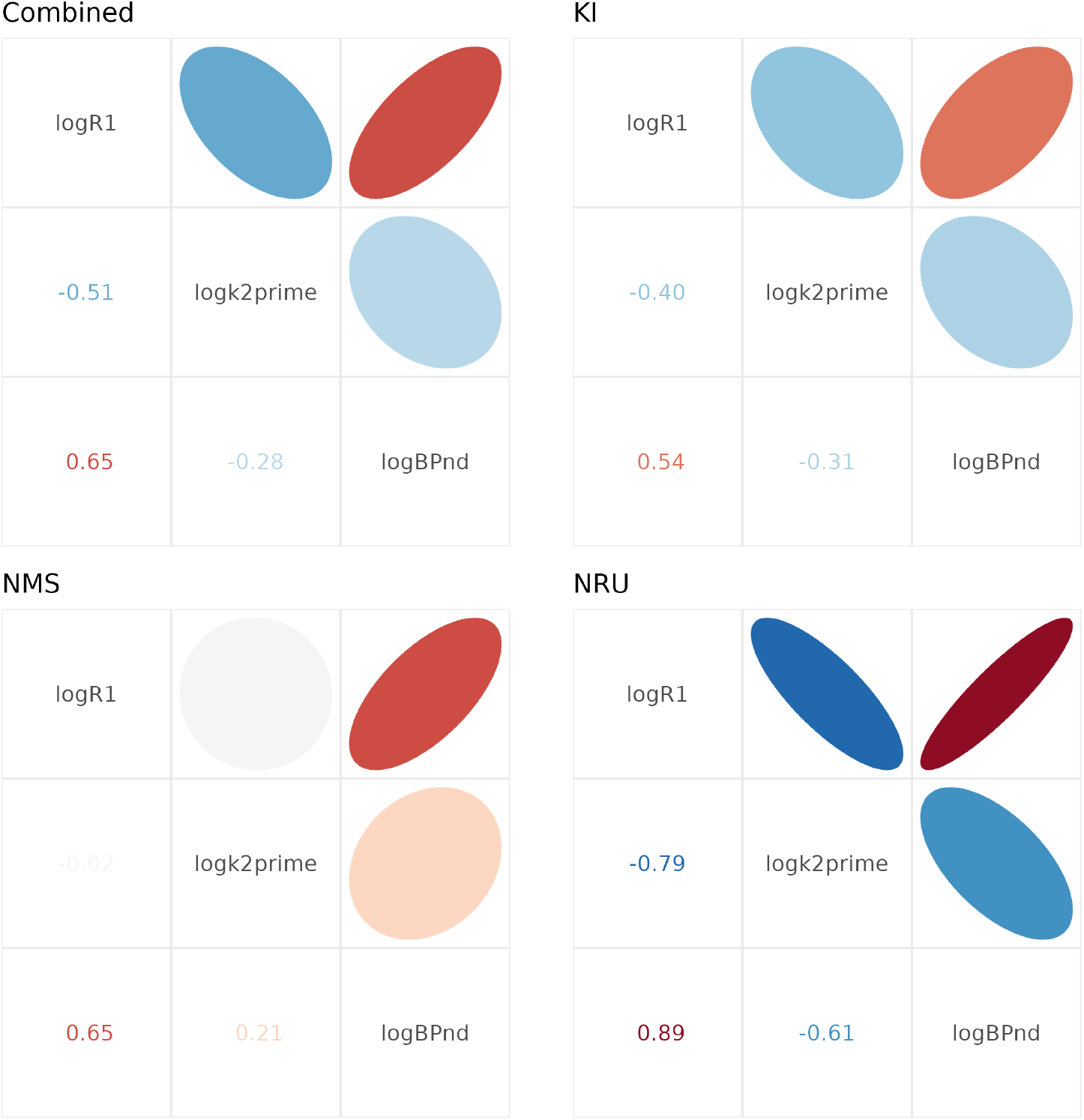
Correlation matrices for individual level deviations for each model.

**Figure S6.**
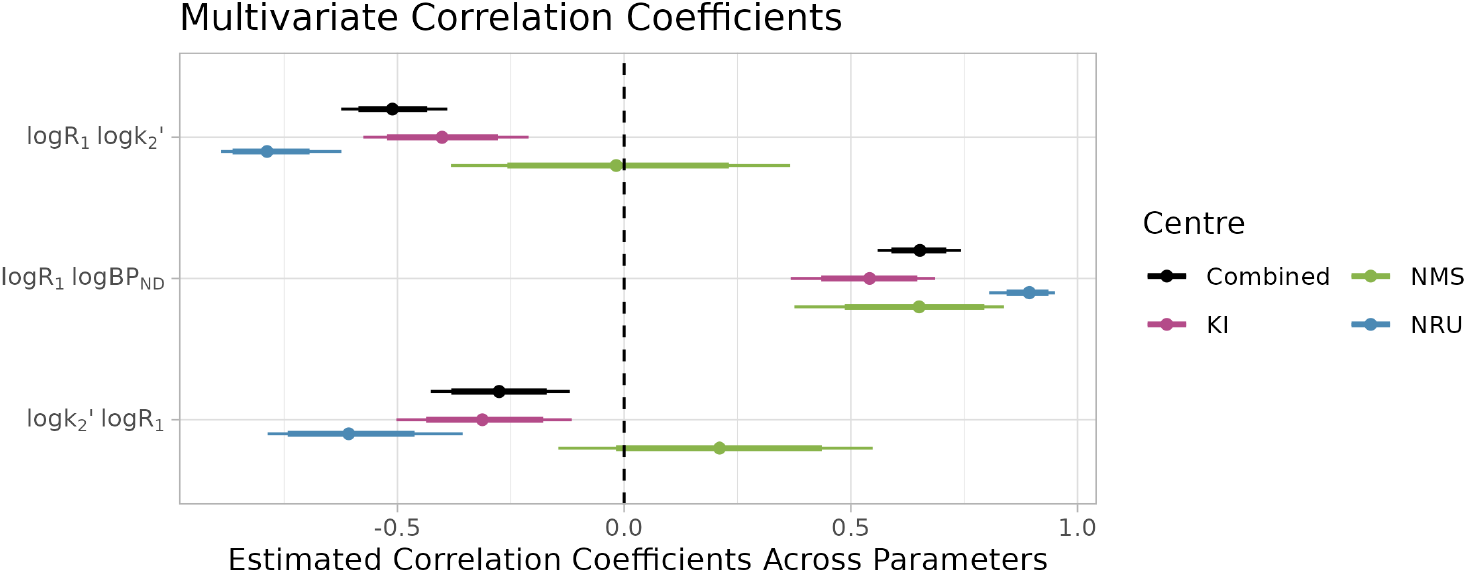
Correlation matrix estimates for individual level deviations with 95% credible intervals.

